# Platelet transcriptome yields progressive markers in chronic myeloproliferative neoplasms and identifies putative targets of therapy

**DOI:** 10.1101/2021.03.12.435190

**Authors:** Zhu Shen, Wenfei Du, Cecelia Perkins, Lenn Fechter, Vanita Natu, Holden Maecker, Jesse Rowley, Jason Gotlib, James Zehnder, Anandi Krishnan

**Affiliations:** Department of Statistics, Stanford University, Stanford, CA; Stanford Cancer Institute, Stanford University School of Medicine, Stanford, CA; Stanford Functional Genomics Facility, Stanford University School of Medicine, Stanford, CA; Department of Microbiology and Immunology, Stanford University School of Medicine, Stanford, CA; Department of Internal Medicine, University of Utah, Salt Lake City, UT; Department of Medicine, Stanford University School of Medicine, Stanford, CA; Department of Pathology, Stanford University, Stanford, CA

## Abstract

Predicting disease progression remains a particularly challenging endeavor in chronic degenerative disorders and cancer, thus limiting early detection, risk stratification, and preventive interventions. Here, profiling the spectrum of chronic myeloproliferative neoplasms (MPNs) as a model, we identify the blood platelet transcriptome as a proxy for highly sensitive progression biomarkers that also enables prediction of advanced disease via machine learning algorithms. Using RNA sequencing (RNA-seq), we derive disease-relevant gene expression in purified platelets from 120 peripheral blood samples constituting two time-separated cohorts of patients diagnosed with one of three MPN subtypes at sample acquisition – essential thrombocythemia, ET (n=24), polycythemia vera, PV (n=33), and primary or post ET/PV secondary myelofibrosis, MF (n=42), and healthy donors (n=21). The MPN platelet transcriptome reveals an incremental molecular reprogramming that is independent of patient driver mutation status or therapy and discriminates each clinical phenotype. Leveraging this dataset that shows a characteristic progressive expression gradient across MPN, we develop a machine learning model (Lasso-penalized regression) and predict advanced subtype MF at high accuracy and under two conditions of external validation: i) temporal: our two Stanford cohorts, AUC-ROC of 0.96; and ii) geographical: independently published data of an additional n=25 MF and n=46 healthy donors, AUC-ROC of 0.97). Lasso-derived signatures offer a robust core set of < 5 MPN transcriptome markers that are progressive in expression. Mechanistic insights from our data highlight impaired protein homeostasis as a prominent driver of MPN evolution, with persistent integrated stress response. We also identify JAK inhibitor-specific signatures and other interferon, proliferation, and proteostasis-associated markers as putative targets for MPN-directed therapy. Our platelet transcriptome snapshot of chronic MPNs demonstrates a proof of principle for disease risk stratification and progression beyond genetic data alone, with potential utility in other progressive disorders.

**Highlights:** Leveraging two independent and mutually validating MPN patient cohorts, we identify progressive transcriptomic markers that also enable externally validated prediction in MPNs.

Our platelet RNA-Seq data identifies impaired protein homeostasis as prominent in MPN progression and offers putative targets of therapy.

**VISUAL ABSTRACT:** 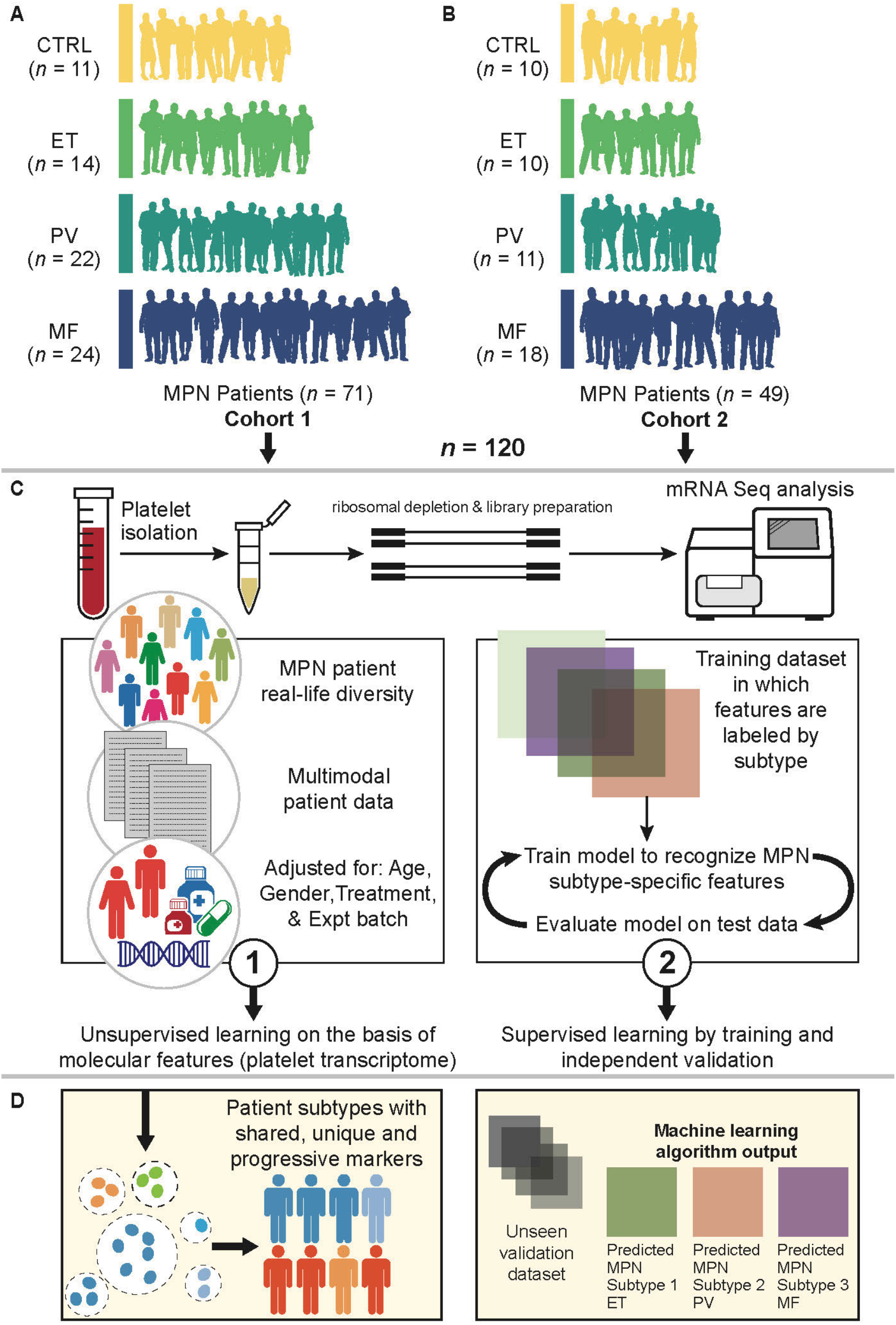

## Introduction

The classic Philadelphia chromosome-negative (Ph^-^) MPNs,(Barbui and Falanga, 2016; Finazzi et al., 2013; Spivak, 2017; Spivak et al., 2014) are clonal disorders of the bone marrow that comprise three clinical phenotypes– essential thrombocythemia (ET), polycythemia vera (PV), and myelofibrosis (MF). These myeloid neoplasms are defined by a combination of morphologic, clinical, laboratory, and cytogenetic/molecular genetic features. The existing genetic landscape (Rumi and Cazzola, 2017; Vainchenker and Kralovics, 2017; Zoi and Cross, 2017) of MPNs primarily involves mutations in three driver genes that lead to constitutive JAK-STAT signaling (*JAK2, CALR, MPL*). Several additional non-driver mutations (see references (Hinds et al., 2016; Nguyen and Gotlib, 2012; Oh and Gotlib, 2010; Rampal et al., 2014; Rumi and Cazzola, 2017; Vainchenker and Kralovics, 2017; Zoi and Cross, 2017) for details) as well as cytogenetic (Spivak, 2017) and epigenetic (Mascarenhas et al., 2011; Tefferi et al., 2011) abnormalities also contribute to disease initiation and progression, and impact both overall survival and potential for progression to acute myeloid leukemia (AML) (Papaemmanuil et al., 2016). Depending on the MPN and stage of disease, these patients may exhibit debilitating constitutional symptoms such as fatigue, pruritus, night sweats, and weight loss; thrombohemorrhagic diathesis and extramedullary hematopoiesis; and an increased propensity for transformation to AML. Although an increase in one or more blood cell lineages contributes to these morbid sequelae, the qualitative abnormalities of myeloid cells that increase vascular risk or disease progression are not well understood. Taken together, a limited understanding exists regarding how genotypic variability contributes to diverse phenotypic presentations and disease natural histories. We were motivated by the current clinical need (Rumi and Cazzola, 2017; Spivak, 2017; Vainchenker and Kralovics, 2017) and the potential for deeper integration of clinical features and genetics with gene expression profiling to improve stratification and management of chronic blood disorders, such as MPNs.

Blood platelets play critical roles in multiple processes and diseases (Rondina et al., 2013; Weyrich, 2014; Weyrich and Zimmerman, 2013), from their traditional function in hemostasis and wound healing to inflammation, immunity, cancer metastasis, and angiogenesis. Platelets originate from bone marrow precursor megakaryocytes as anucleate fragments with a distinctive discoid shape. Platelets contain a complex transcriptional landscape of messenger RNAs (mRNAs), unspliced pre-mRNAs, rRNAs, tRNAs and microRNAs (Rowley et al., 2012; Schubert et al., 2014; Simon et al., 2014; Weyrich, 2014). Most platelet RNA expression results from the transcription of nuclear DNA in the megakaryocyte (Davizon-Castillo et al., 2020; Rowley et al., 2012), and thus reflects the status of the megakaryocyte at the time of platelet release into the circulation, as well as subsequent splicing events, and selective packaging and inter-cellular RNA transfer. There is emerging evidence (Best et al., 2015; Campbell et al., 2018; Clancy and Freedman, 2016; Cunin et al., 2019; Middleton et al., 2019) that the molecular signature of platelets may be changed in disease conditions where these processes are altered, including via inter-cellular transfer (Clancy and Freedman, 2016; Cunin et al., 2019) of cytosolic RNA. In the context of MPNs, the platelet transcriptome therefore not only represents a critical biomarker of megakaryocytic activity (Gilles et al., 2012; Krause and Crispino, 2013; Wen et al., 2016; Wen et al., 2015; Woods et al., 2019), but also provides a snapshot of the underlying hemostatic, thrombotic, and inflammatory derangements associated with these hematologic neoplasms and the potential impact of treatment (Woods et al., 2019).

To date, no one composite study (Krishnan et al., 2017) (Gangaraju et al., 2020; Guo et al., 2019; Rampal et al., 2014; Schischlik et al., 2019; Skov et al., 2011; Skov et al., 2012) has evaluated the disease-relevant platelet transcriptome in a sizeable clinical cohort of all three subtypes of Ph^-^ MPNs. Here, we extend our prior preliminary work(Krishnan et al., 2017) toward a comprehensive analysis of disease-relevant (Cummings et al., 2017) platelet RNA-sequencing (RNA-seq) in two temporally independent and mutually validating cohorts of all three MPN subtypes, ET, PV, and MF (primary or post ET/PV secondary). We demonstrate marked differences in platelet gene expression across the MPN spectrum, which also permits robust validated (temporal and geographical) prediction of MF. In addition to identifying novel gene expression signatures impacted by the JAK1/JAK2 inhibitor ruxolitinib (RUX), platelet profiling reveals MPN-altered pathways that may be targets for future drug development.

## Results

### Two independent MPN clinical cohorts and closely replicated platelet transcriptome

We prepared highly purified leukocyte-depleted (Amisten, 2012; Rowley et al., 2011) platelets from peripheral blood samples of two cohorts (approximately 2 years apart; Stanford single-center) of patients with a diagnosis of MPN (including provisional) at the time of sample acquisition, and included healthy controls in each cohort as reference (cohort 1, n = 71, and cohort 2, n=49; **Figure 1**, **Table S1A-B**). Only 2 patients (2%) received a change in diagnosis from MPN (MF) to a non-MPN phenotype; and were therefore excluded from all downstream analyses (**Figure 2A** principal component analysis plot panel-3 identifies these 2 outliers). Our two-cohort study was specifically designed (before knowledge of other subsequent studies) for the explicit purpose of validation, not only of inter-cohort RNA-seq results (Kukurba and Montgomery, 2015) but also to evaluate temporal validation (Moons et al., 2012) of our prediction model (see Methods, **Figure 5C,E,F**). **Figure S1** demonstrates our established(Rowley et al., 2011; Rowley et al., 2019) high-quality and highly efficient experimental framework toward a rigorous platelet RNA-seq approach. Clinical features of the MPN patients are shown in **Figure 1**; and listed in **Table S1A-B**. The distribution of key variables was closely matched between the two cohorts by MPN subtype (**Figure 1A**), age (**B**), gender (**C**), *JAK2/CALR* mutation status (**D**) and treatment (**E**). Any inter-patient variability in patient age, gender and treatment were adjusted as confounding factors in all downstream gene expression analyses (see Methods). Clinical laboratory measures (**Figure 1F**) at the time of sampling reflected the phenotype of the MPN subtypes (please see figure legend for detailed statistical comparisons). The two cohorts of platelet transcriptome data (**Figure 1G**, normalized transcript counts) adjusted for patient age, gender and all treatment as confounding factors were also highly correlated (R^2^ = 0.89), thus demonstrating high inter-cohort validation of gene expression that then enabled us to combine our two RNA-seq datasets into a final integrated data set of enhanced statistical power for downstream analyses, especially prediction modeling (**Figure 5**). Together, this platelet transcriptome compendium comprises 118 human peripheral blood samples from healthy controls (n=21) and World Health Organization-defined MPN patients (24 ET, 33 PV and 40 MF) that include seven untreated, and 92 either on cytoreductives/biologics (*e.g.* ruxolitinib, hydroxyurea, interferon-alpha), anti-thrombotic agents (e.g. aspirin, warfarin), or a combination and captures the real-life diversity among MPN patients. Our cross-sectional design here capturing patients from all three MPN subtypes also serves as a practical alternative to the longitudinal approach, though ideal, is likely difficult to implement in these chronic disorders with often decades of disease.

**Figure 1:**
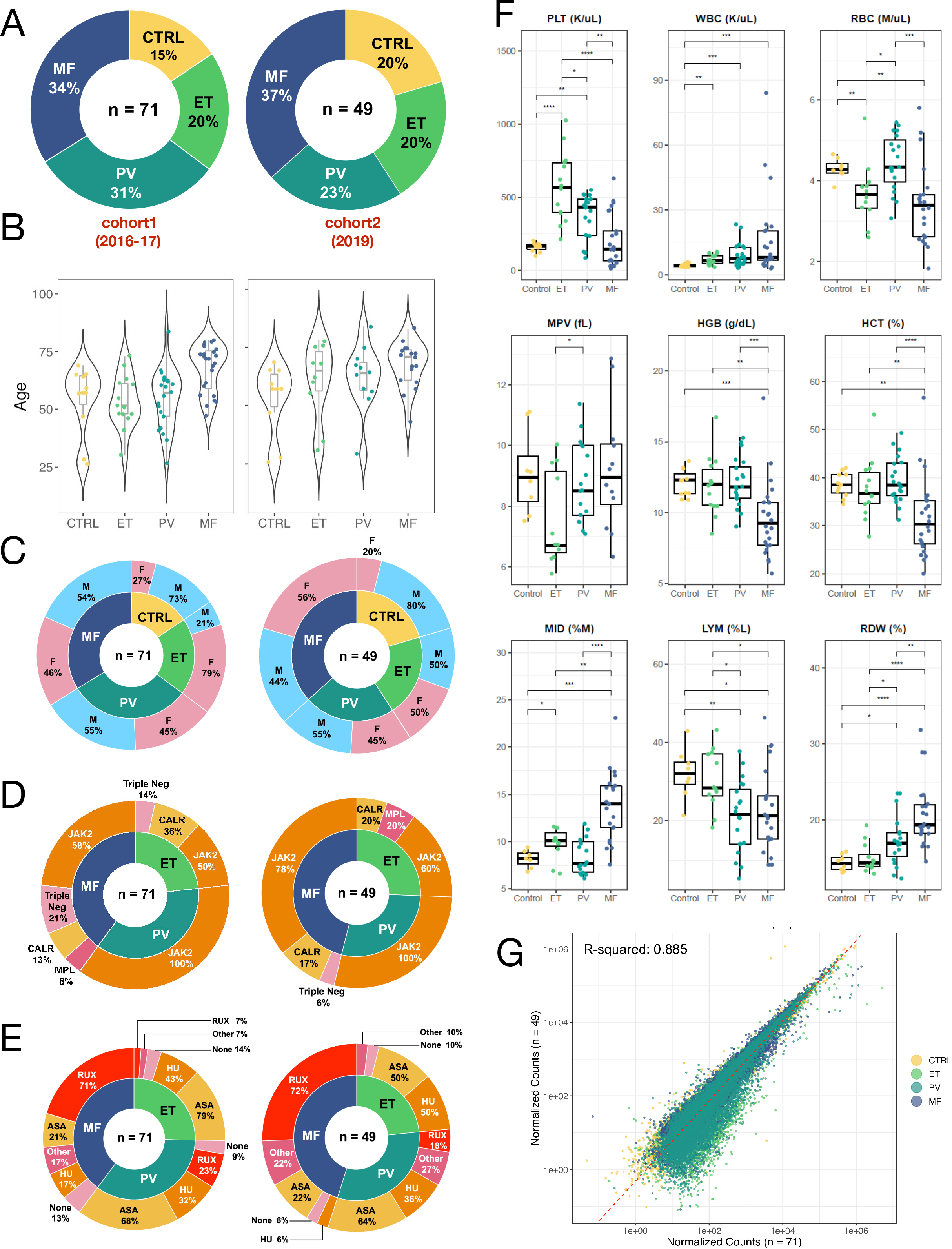
Two independent MPN clinical cohorts and closely replicated patient variables. **A**, Similarity in distribution of MPN subtypes between two cohorts of MPN patients (Stanford single-center; approximately 2 years apart: cohort 1: 2016-17, n=71; and cohort 2: 2019, n=49); the majority subtype is MF in both cohorts (34% of n=71 and 37% of n=49). **B**, Comparable distribution of age across MPN subtypes in the two cohorts. Violin plots of patient age from each MPN subtype reflect clinical expectation, with a fairly identical match between the two cohorts. Slightly higher median age noted in the second cohort for ET and PV patients alone. **C,** Comparable and balanced distribution of gender across MPN subtypes in the two cohorts. Larger percentage of male healthy controls in both cohorts and smaller percentage of males in ET cohort 1 noted. **D,** Matched distribution of primarily *JAK2* and *CALR* mutational status across MPN subtypes in the two cohorts. JAK2 is the most common mutation across all three subtypes; 100% of PV and over 50% of ET and MF patients have JAK2 mutation in both cohorts. Mismatch between cohorts on the MPL/triple negative patients is noted as a natural consequence of the rarer prevalence of these mutations; and therefore, not the primary focus of this work. **E,** Diversity of MPN patient therapies across the two cohorts reflecting current clinical practice. The majority being aspirin (ASA) in ET/PV patients and the *JAK*-inhibitor, ruxolitinib in MF. Note that a given patient may be on more than one treatment and therefore, the total treatment percentage in this graphic may not equal 100. To control for any inter-patient variability, all treatment, in addition to patient age and gender are adjusted as confounding factors in downstream gene expression analyses. **F,** Representative clinical laboratory variables, as box plots, measured at the same date and time as platelet sampling. Compared to controls, MPN patients show larger variance (inter-quartile range, IQR), and reflect current clinical knowledge. Groups differ primarily only with respect to blood cell counts (platelet/RBC/WBC); and show the greatest differential in MF. Note higher platelet count in ET with a concomitant lower mean platelet volume, higher red blood cell count in PV and lower red blood cell count in MF with concomitantly lower hemoglobin, hematocrit and higher red cell width. Wilcoxson signed rank tests marked by asterisks indicate a statistically significant difference between any two groups (* p<=0.05; ** p<=0.01; *** p<=0.001; **** p<=0.0001). **G,** High correlation (R^2^ = 0.89) of platelet gene expression as assessed by normalized counts of matched transcripts in each cohort between each of controls, ET, PV and MF. The two-cohort collective sample size totals n=120, affording increased statistical power for subsequent analyses.

**Figure 2:**
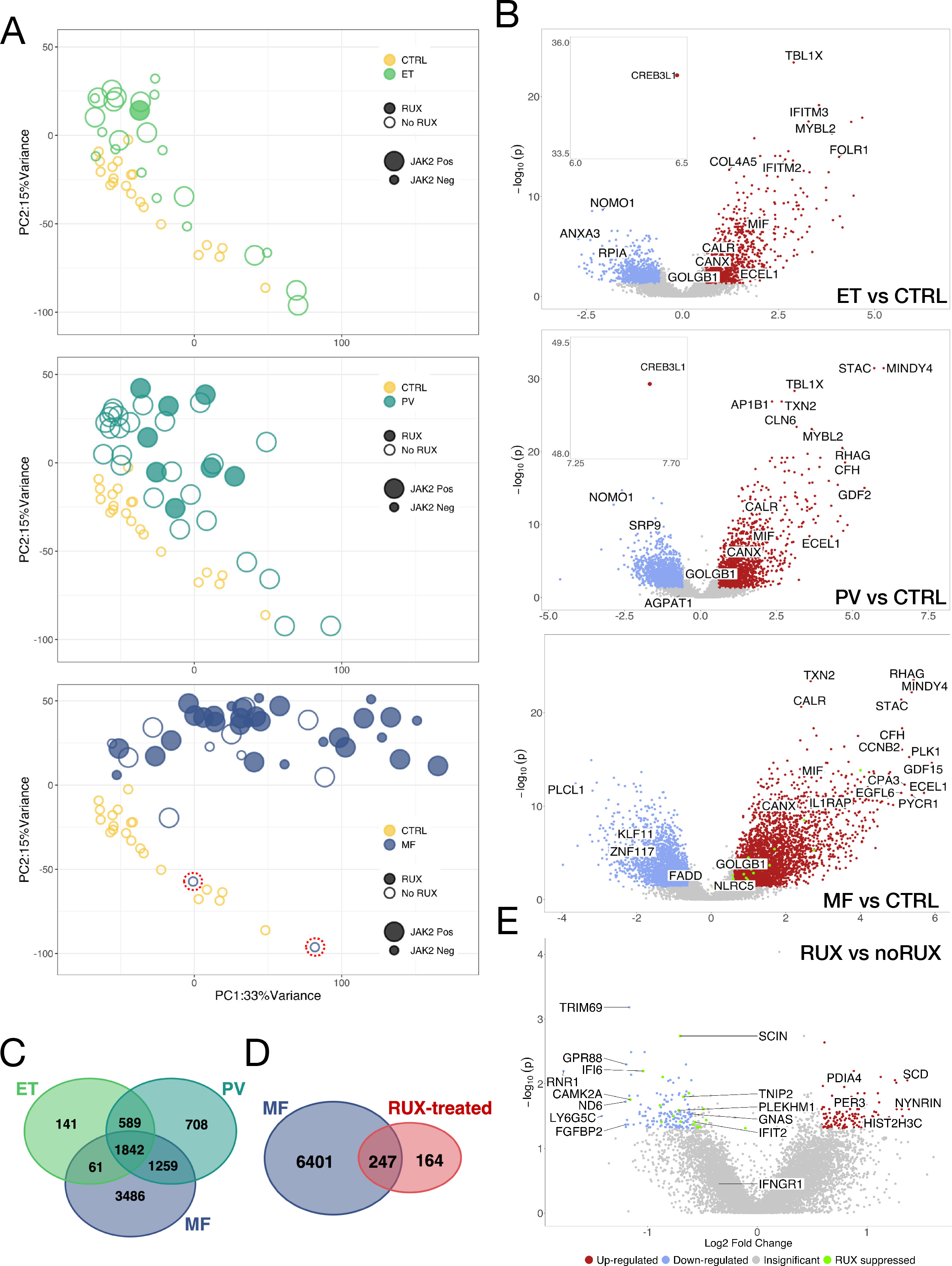
MPN platelet transcriptome distinguishes disease phenotype and reveals phenotype- and JAK-inhibitor specific signatures. **A**, Unsupervised principal component analysis (PCA) of normalized platelet gene expression counts adjusted for age, gender, treatment and experimental batch. Three panels of PC1 and PC2 colored by MPN subtype; and each contrasted with controls (n=21, yellow): ET (n=24, top, light green), PV (n=33, middle, dark green) and MF (n=42, bottom, dark blue). Circles filled or open mark presence or absence of ruxolitinib treatment; and size of circles, smaller or larger, indicate presence or absence of *JAK2* mutation. The first two principal components account for 48% of total variance in the data. **B,** Volcano plots (three panels as **a** above of ET, PV and MF) of differential gene expression showing statistical significance (negative log10 of p-values) versus log2 fold change of each gene. Significant up-regulated and down-regulated genes are those with p-values (FDR) smaller or equal to 0.05 and absolute value of fold changes larger or equal to 1.5**. C,D,** Venn Diagram comparisons of MPN differential gene expression lists. In **c,** each of ET, PV and MF is contrasted with controls; identifying gene sets that are shared (n=1504, FDR < 0.05) as well as unique to each subtype. In **D,** differential in the RUX-treated cohort is contrasted with MF vs controls. Differential in gene expression in RUX-treated cohort is a fraction of the total differential noted in the MF transcriptome. **E,** Volcano plot of differential gene expression between MPN patients treated with ruxolitinib and not. A small subset of overlapping differential genes that are upregulated in MF (**B,** bottom panel) and suppressed in the RUX-treated cohort (**E**) are colored in green.

### MPN platelet transcriptome distinguishes disease phenotype and reveals phenotype- and JAK-inhibitor specific signatures

Given the phenotypic overlap, yet also differences in disease behavior and prognosis between ET, PV and MF, we hypothesized that there may be MPN subtype-specific differences in gene expression that are independent of *JAK2/CALR/MPL* mutation status. In addition, we hypothesized that MPN platelets are likely to be enriched for subtype-specific biomarkers that may otherwise be missed in blood/plasma/serum sources (Schischlik et al., 2019; Skov et al., 2011; Skov et al., 2012). Therefore, we compared platelet transcriptomic expression in each of the three MPN subtypes with that of controls and discovered a shared gene set that is also progressively differentiated across the MPN spectrum (ET/PV to MF **Figure 2 A-C**). First, unsupervised principal component analysis (PCA) of MPN patients and controls data (**Figure 2A**) confirmed that the collective variability from the first two principal components (accounting for 48% of total variance), after adjusting for age, gender, treatment and experimental batch, was MPN disease status, with increasing differentiation by subtype. Next, differential gene expression analysis (DGEA, volcano plot, **Figure 2B, C**) efficiently distinguished each of the MPN subtypes and resulted in highly significant expression signatures (adjusted p-value/FDR <0.05) with 2634 genes differentially regulated in ET (1364 up and 1269 down), 4398 in PV (2098 up and 2300 down), and 6648 in MF (3965 up and 2683 down). A subset of 100+ long non-coding RNAs and pseudogenes also constituted the significant (FDR<0.05) differential expression across MPNs (**Table S2A-C**).

Specifically, DGEA also uncovered shared and unique genes between all three MPN phenotypes, thus offering a potential core set of genes involved in MPN pathogenesis (**Figure 2C**, with associated heatmaps in Figure 3 and pathways in Figure 4). The shared gene set at FDR < 0.05 constituted 654 up-regulated genes, with a predominance of molecular pathways involving *myeloid cell activation in immune response and membrane protein proteolysis*; and 361 down-regulated genes, reflecting negative regulation of hematopoeisis and negative regulation of transmembrane receptor protein serine/threonine kinase signaling as a consistent pathogenetic mechanism. The upregulated genes belonged to the endoplasmic reticulum/ER-Golgi intermediate compartment, and included a particularly high expression of the cAMP-response element binding transcription factor, *CREB3L1*(Sampieri et al., 2019) implicated in cell differentiation and inhibition of cell proliferation, with concomitant high expression of ER chaperones(Clark et al., 2002) calreticulin (*CALR*), calnexin (*CANX*); transport *factors:* golgin *(GOLGB1)* and folate receptor *FOLR1.* Platelet alpha granule proteins *(F5, VWF, MMP14),* several collagens (*COL10A1, COL18A1, COL6A3*), immune/inflammatory (*IFITM2/3/10, FCGR2A, TMEM179B*), *Cathepsins (C/Z/D, MIF, PTGES2*) and proliferation mediator genes (*CDK1, CCNG1, BMP9/GDF2, LAPTM4B, PSENEN*) also constituted the MPN shared set. Downregulated genes, on the other hand, were predominantly within transcription factor complexes, and included opposing expression of *CREB1* (vis-à-vis *CREB3L1* above), calcium-calmodulin protein kinases, *CAMK4*, *SMAD1* and β-catenin *CTNNB1* together pointing to dysregulated calcium (Ca^2+^) homeostasis in the ER lumen.

**Figure 3:**
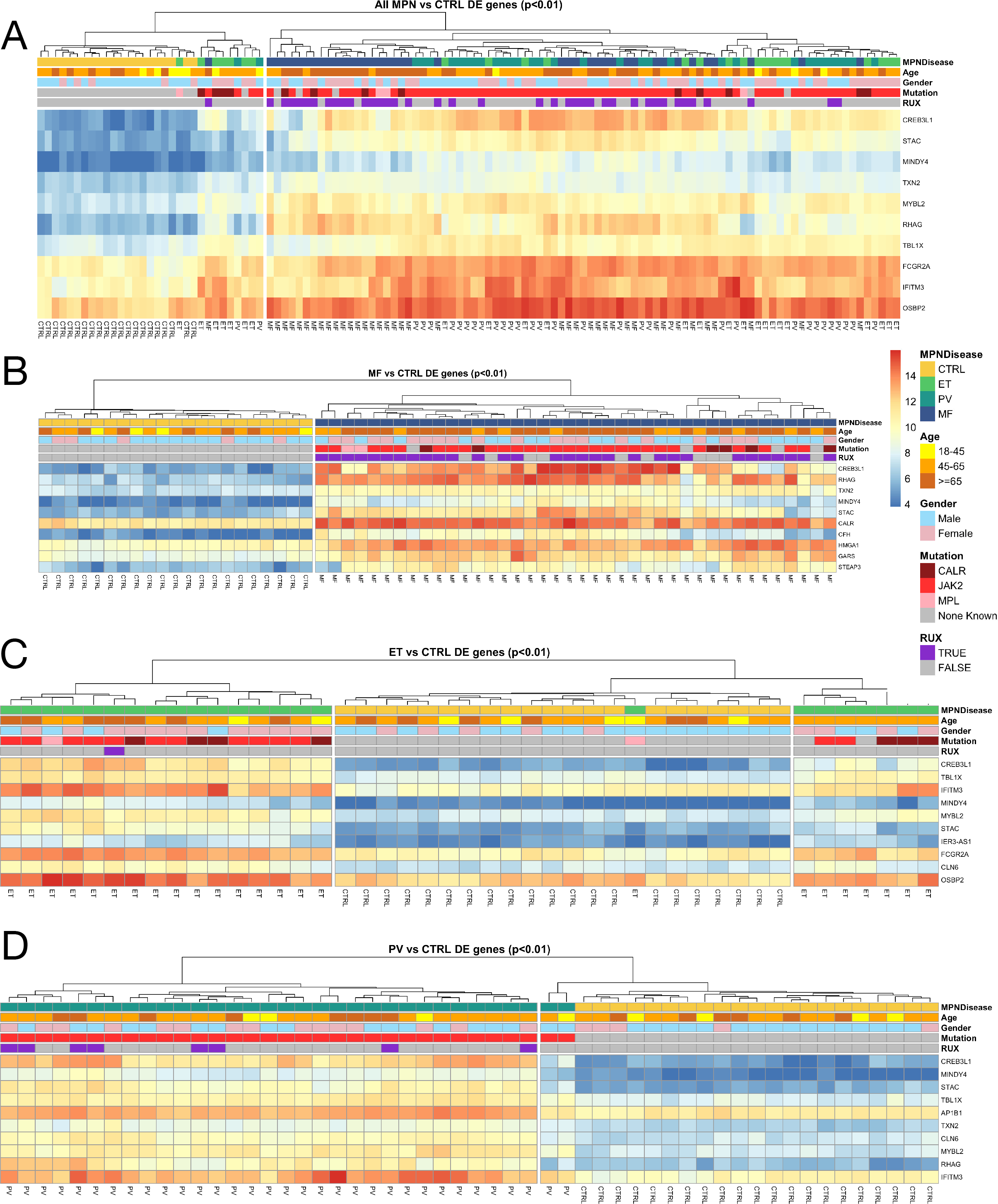
Graded differential expression by MPN phenotype and driver mutation status. **A-D**, Hierarchically clustered heatmaps of the top 10 differentially expressed genes (DEGs) from controls (FDR<0.01) of all MPN (**A**), and MF, ET, and PV each separately (**B-D)**. Colored annotation is provided to indicate MPN subtype, age, gender, mutation and ruxolitinib treatment. Rows indicate gradation in gene expression on a blue (low) to red (high) scale. Columns indicate sample type from controls (CTRL) to ET, PV and MF.

**Figure 4:**
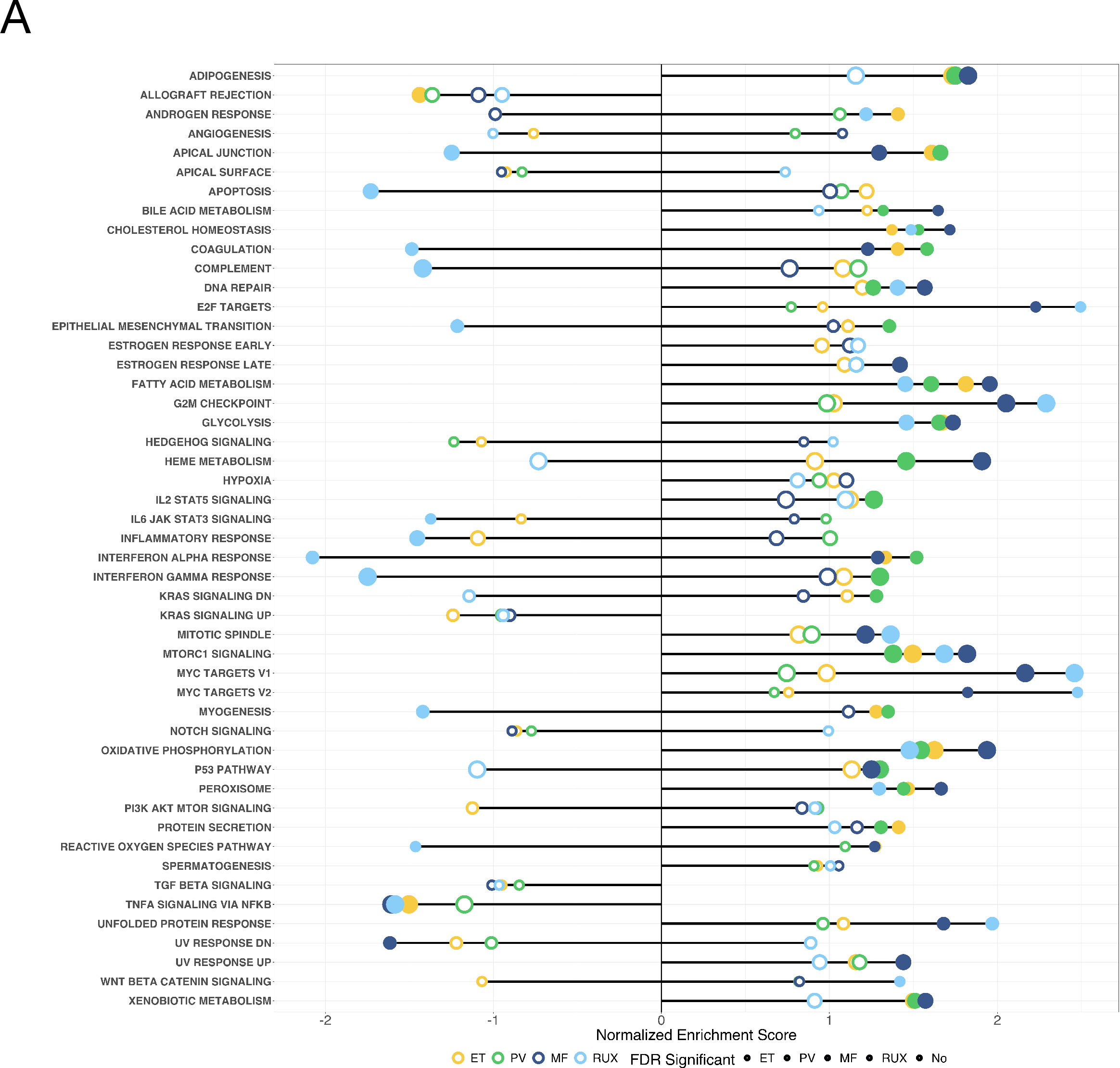
Altered immune, metabolic, and proteostatic pathways underlie each MPN phenotype. **A**, Pathway-enrichment analysis of genes with MPN subtype-specific expression (color indicated; light green ET, dark green PV, and dark blue MF) overlayed with ruxolitinib-specific expression (light blue). Each point represents a pathway; the *x*-axis gives the normalized enrichment score, which reflects the degree to which each pathway is over or under-represented at the top or bottom of the ranked list of differentially expressed genes, normalized to account for differences in gene set size and in correlations between gene sets and the expression data set. The y-axis lists the detail-level node of the most enriched pathways; solid lines mark GSEA-recommended(Subramanian et al., 2005) Bonferroni-corrected statistical significance criterion of FDR < 0.25 for exploratory analyses. Dotted lines mark FDR > 0.25 and therefore, not sufficiently significant, yet visualized alongside solid lines to retain overall context (upper-level parent nodes of the detail-level pathways are provided in **Table S3A-C**). Multiple immune and inflammatory pathways are consistently significantly enriched across ET, PV and MF (and suppressed in the ruxolitinib-treated cohort). MF is differentiated from ET and PV through dysregulation of additional molecular processes for cellular development, proliferation, metabolism and DNA damage.

**Figure 5:**
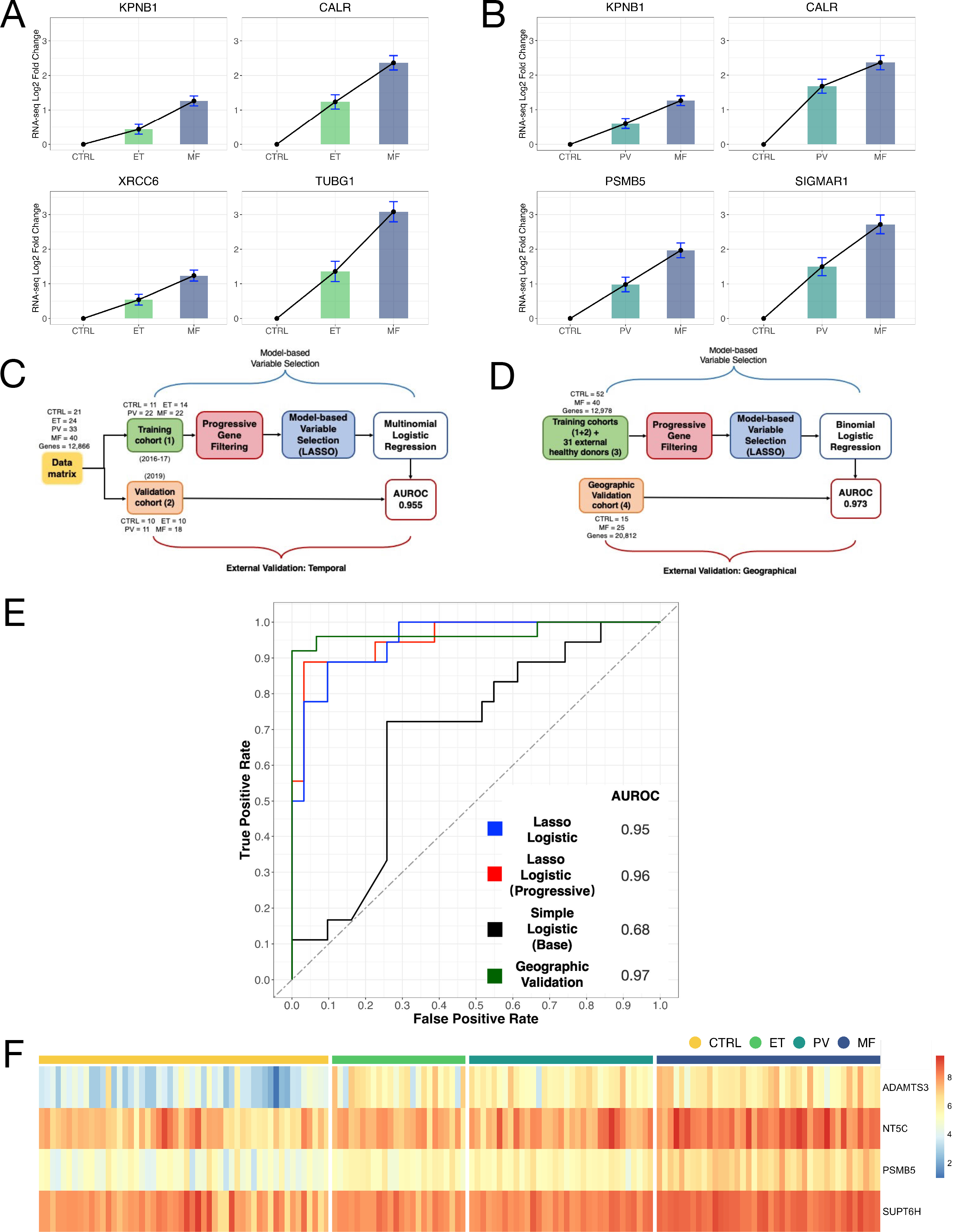
Prediction of MF based on unique and progressive MPN platelet transcriptome. **A, B** Top few genes (out of 3000+) demonstrating monotonic progressive gene expression (log 2 fold change in expression *y*-axis, FDR < 0.01, *Mann-Kendall* test with Bonferroni correction) across *x*-axes **A**, CTRL-to-ET-to-MF and **B**, CTRL-to-PV-to-MF. **C,** Lasso-penalized multinomial logistic regression model under temporal validation *i.e.* trained on Stanford cohort 1 (n=71, 2016-17, Figure 1a) and validated on Stanford cohort 2 (n=49, 2019, Figure 1a) as test set. **D,** Lasso-penalized multinomial logistic regression model under geographical validation using two independently published datasets for training (cohort 3, n=31 healthy controls in addition to Stanford 1 & 2 cohorts) and validation (cohort 4, n=25 MF and n=15 healthy controls). **E,** Receiver Operating Curves (ROC) toward MF prediction under conditions of temporal (**C**) and geographical (**D**) validation. Temporal validation uses three layered models: i) baseline, with no gene expression data but patient age, gender and mutation alone; ii) adding entire MPN platelet transcriptome and iii) adding MPN progressive genes alone. Outperformance of the progressive transcriptome model (red curve, AUROC=0.96) vis-a-vis the entire transcriptome dataset (blue curve, AUROC=0.95) and lastly, the baseline model without gene expression (black curve, AUROC=0.68). Geographical validation using the progressive transcriptome model also demonstrates independent high MF predictive accuracy (green curve, AUROC=0.97). **F,** Heatmap of top recurring Lasso-selected progressive genes for each of controls (left column, CTRL, yellow bar), ET (light green), PV (dark green) and MF (dark blue). Rows indicate gradation in gene expression on a blue (low) to red (high) scale. Columns indicate sample type (CTRL, ET, PV and MF).

Differential markers in each of ET, PV and MF also highlight candidate genes as potential mediators of the pro-thrombotic and pro-fibrotic phenotypes in MPNs. In ET and PV, a strong thromboinflammatory profile (Barbui and Falanga, 2016; Gangaraju et al., 2020) is revealed by the upregulation of several interferon inducible transmembrane genes (*IFITM2, IFITM3, IFITM10, IFIT3, IFI6, IFI27L1, IFI27L2*), interleukin receptor accessory kinases/proteins (*IRAK1, IL15, IL1RAP, IL17RC) and* several solute carrier family genes (*SLC16A1, SLC25A1, SLC26A8, SLC2A9)* as glucose and other metabolic transport proteins, and coagulation factor V (*F5)*. In MF, fibrosis-specific markers were identified by an additional focused comparison of MF patients versus ET and PV (**Figure S2**), showing increased expression of several pro-fibrotic growth factors (*FGFR1, FGFR3, FGFRL1*), matrix metalloproteinases (*MMP8, MMP14*), vascular endothelial growth factor A (*VEGFA*), insulin growth factor binding protein (*IGFBP7*), and cell cycle regulators (*CCND1, CCNA2, CCNB2, CCNF*). GSEA of this MF-focused comparison again highlighted potential underlying molecular dysregulation in MPNs that likely contribute to the fibrotic phenotype (*e.g.* unfolded protein response, mTORc1 signaling, MYC/E2F targets, oxidative phosphorylation, **Figure 4**). Having defined differential gene expression signatures by MPN subtype, we then explored how platelet gene expression profiles differed in patients by treatment, focusing for this work on the JAK1/JAK2 inhibitor, ruxolitinib (RUX, **Figure 2E**). DGEA on the platelet transcriptome in RUX-treated patients identified over 400 core significant genes changed in response to treatment (**Figure 2E and Supplementary Tables S4 and S5**). At-least two-fold reduction in expression was noted in genes associated with interferon-stimulation (*IFI6, TRIM69, LY6G5C*), platelet-mediated apoptosis, *FASLG*, G-protein coupled receptor, *GPR88,* calcium-calmodulin protein kinase, *CAMK2A* and fibroblast growth factor binding protein, FGFBP2, followed by over 150 genes with at-least one-fold reduction in expression in the RUX-treated cohort (**Supplementary Table S5**). These included genes downregulated within classical pathways of adaptive immune response (*TNFSF13B, B2M, HMGB1*), response to oxidative stress (*COX1, COX2, COX15, TP53*) and myeloid activation (*TNIP2, FTH1, TOLLIP, RAB6A*).

In addition to confirming previous observations (Arcaini and Cazzola, 2018; Kleppe et al., 2018; Spivak, 2017; Vainchenker et al., 2018) on the anti-inflammatory and immunosuppressive effects of *JAK* inhibition by RUX (e.g. downregulation in our RUX-treated cohort of *IL1RAP, CXCR5, CPNE3, ILF3*), we identified new gene clusters responsive to RUX in the inhibition of type I interferon (e.g. *IFIT1, IFIT2, IFI6*), chromatin regulation (*HIST2H3A/C, HIST1H2BK, H2AFY, SMARCA4, SMARCC2*), and epigenetic methylation in mitochondrial genes (*ATP6, ATP8, ND1-6 and NDUFA5*). Recent literature probing the mechanisms of action of ruxolitinib in other disease settings, including SARS-CoV-2 (Yan et al., 2021; Zhou et al., 2014) confirm our observations in MPNs.

A direct comparison restricted to only differentially expressed genes (FDR<0.05) in *RUX-treated vs not* with *MF vs healthy controls* revealed less than 5% overlap (**Figure 2D**), reflecting potentially the extent of the impact of treatment by ruxolitinib relative to the substantive disease burden in MF. Focusing further on the directionality of the changes observed, we found just 18 genes that were increased in MF and suppressed in the RUX-treated cohort (**Figure 2E**, colored green); and 9 vice versa (**Table S3A-B**). Despite the small overlap, we capitalized on the converging genes to better define molecular and physiological pathways underlying the effect of RUX in MPNs. The 18 RUX-down-regulated genes followed expected mapping to immunosuppression through interferon- and cytokine-mediated signaling pathways. The 9 genes that were upregulated with RUX in MF mapped to previously undescribed effect of the drug in select G-protein-coupled receptor and chemokine activity (*e.g. CXCR5, GPR128/ADGRG7*), semaphorin signaling (*SEMA3C*) and circadian regulation (*PER3*). A sub-cohort analysis of RUX-treated and RUX-naïve MF patients alone also identified downregulation of interferon-stimulated genes (*e.g. SLFN12L),* G-protein-coupled receptors *(S1PR5), f*ibroblast growth factor binding protein *(FGFBP2) and* tyrosine kinase *(LCK)*.

### Graded differential expression by MPN phenotype and driver mutation status

Unsupervised hierarchical clustering was used to more precisely define the nature of MPN platelet transcriptome variability from controls, as well as between and within MPN subtypes. **Figure 3** reveals a spectrum of expression in the MPN platelet transcriptomic profile using just the top 10 highly significant differentially expressed genes by disease status: (a) all MPN vs controls, (b) MF vs controls, and (c, and d) ET- and PV-vs controls. As shown in **Figure 3A**, all MPN patients clustered into two distinct groups: a larger group of 87 ET, PV and MF patients clustered independently from the 21 controls, whereas a smaller group of 10 ET, PV and MF patients formed a homogeneous cluster of their own closer to controls reflecting a varying gradation with respect to the top-10 gene expression by MPN subtype (patient variables annotated above the heatmaps offer additional context, particularly on mutation status and RUX therapy). In the larger cluster, while we observed a graded overlap in platelet RNA signatures between ET and PV, a more distinct expression pattern characterized the more advanced population of MF patients (*PC1 correlates with MF disease risk by the Dynamic International Prognostic Scoring System (Passamonti et al., 2010), DIPSS*, **Figure S3**). These data collectively highlight the importance of phenotype-modifying genes that are independent of JAK2*/CALR/MPL* mutation status.

Untangling other mechanisms beyond the few genes that are recurrently mutated is critical for defining subtype-specific risk and for identifying molecular pathways for targeted therapy. Accordingly, we sought to refine the molecular classification of MPN by associating platelet gene expression profiles with the corresponding subtype, and yielded a core set of 10 highly-significant preferentially expressed genes for each: (i) MF (**Figure 3B**), defined by high mRNA expression of proteostasis-associated *CREB3L1* and *CALR,* and megakaryocyte-erythroid differentiation stage-associated *RHAG* (Caparros-Perez et al., 2017; Zeddies et al., 2014) ii) ET (**Figure 3C**), marked by comparatively high expression of interferon-related genes *IFITM2*/*3,* immune response *FCGR2A*, and proliferation-associated *STAT5* target *OSBP2*; and iii) PV (**Figure 3D**), marked by overlapping signatures with ET in inflammation-associated *IFITM3* and *TBL1X*, and the B-myb promoter, *MYBL2*; with MF in the maturation-associated *RHAG* at variable expression; and with both ET and MF in the high expression of *CREB3L1*, and cell survival-associated *MINDY4* and *STAC*.

### Altered immune, metabolic, and proteostatic pathways underlie each MPN phenotype

Our analysis of MPN platelet RNA-seq enabled identification of altered MPN pathways that might be amenable to drug therapy. To understand the biological significance of transcriptional changes, we performed pathway-enrichment analysis and identified signaling pathways that are differentially activated between MPN subtypes (**Figure 4**). Gene set enrichment analysis (GSEA, see Methods) of Hallmark gene sets found that MPN (stratified by subtypes, ET, PV and MF) induces genes related to pathways with known immune modulatory functions (**Figure 4**, notably interferon alpha response in ET, PV and MF, and IL2 STAT5 signaling, and interferon gamma response specifically enriched in PV). Moreover, among the most enriched gene sets, MPN pathology induces robust activation of oxidative phosphorylation (OXPHOS) and mTORC1 signaling pathways, with increasing enrichment and significance by MPN subtype (FDR <0.0001 in MF). Pathways of reactive oxygen species (ROS) production paralleled activation of mTORC1 in MF. Other complementary metabolic pathways paralleled OXPHOS activation, with significant enrichment of bile and fatty acid metabolism, cholesterol homeostasis and adipogenesis, most pronounced in MF and variably expressed in ET and PV. Coagulation- and complement-associated gene sets were expectedly enriched across ET, PV and MF. What is particularly noteworthy is that in MF, cell cycle progression and proliferation pathways reveal significant enrichment (FDR <0.001) around c-MYC and E2F targets, and G2M checkpoint pathways; and unfolded protein response emerges as a key factor, likely attributable to ER stress (see *CREB3L1, CALR* overexpression in **Figures 2,3**). Representative GSEA profiles are shown in **Figure 4** and the full list of enriched pathways and gene sets detailed in **Table S4A-C**. The MPN pathways exhibiting significant transcriptional regulation by GSEA are consistent with our observations at the individual gene level for upregulated and downregulated transcripts, specifically those upregulated in MF. Taken together, these data demonstrate that in addition to immune factors such as type I/II interferons and dysregulation of interleukin-dependent inflammatory responses, which have been linked to MPNs, platelet transcriptional signatures of proliferation, metabolic, and proteostasis signaling are a feature of MPN pathogenesis (**Figure S4** captures the relative enrichment by subtype of MPN molecular pathway categories as a concept model of MPN progression).

### Prediction of MF based on shared, unique and progressive MPN platelet transcriptome

Current knowledge of MPN genetic, cytogenetic or epigenetic abnormalities are limited(Spivak, 2017; Spivak et al., 2014) in their ability to enable prediction of disease progression or evolution of a given patient, from ET/PV phenotype to MF. In order to investigate the potential of platelet transcriptomic parameters to enable MF prediction, we constructed LASSO penalized (Tibshirani, 1996) regression classifiers (machine learning R package *glmnet)* to discriminate MPN subtypes from each other, and from healthy controls (**Figure 5A-E**). We apply two rigorous (Moons et al., 2012) external validation conditions **(Figure 5C,D):** i) training and independent temporal validation (**Figure 5C**) leveraging the Stanford two-cohort design and ii) geographical validation (**Figure 5D**) using two independently published platelet transcriptome datasets: first, from Rondina et al., 2020 on an additional n=31 healthy donors integrated with the Stanford datasets as training; and second, from Guo et al., 2020 n=25 MF and n=15 healthy donors as geographical validation of the Lasso algorithm (**Figure 5D**). Our temporal validation constituted three types of models i) baseline (no transcriptome, but age, gender and driver mutation status as reference variable information available not only for patients but also healthy donors); ii) full platelet transcriptome integrated with the above baseline; and iii) subset platelet transcriptome that exhibits progressive differentiation from controls to ET to MF or controls to PV to MF (>3000 genes, top few of each comparison visualized to demonstrate progressive gene expression, **Figure 5A-B**) integrated with the above baseline (progressive subset is selected unbiased as part of the Lasso learning procedure). Comparison of the classification potential among the three models demonstrated that the progressive platelet transcriptome model (**Figure 5E** red curve) substantially outperformed the baseline model (**Figure 5E** black curve) and was slightly better than the full transcriptome model (**Figure 5E** blue vs red curves) in the classification of ET, PV and MF. Predicted probabilities for all three models are shown in **Table S5A-C**. Lasso logistic regression classifiers to predict MF with each of the models under the first temporal two-cohort training-validation split of baseline, full, and progressive transcriptome each achieved area under the receiver-operating characteristic curve (AUROC) of 0.68, 0.95, and 0.96 respectively. Outperformance of the progressive transcriptome model was validated in our subsequent independent external geographic validation (**Figure 5E** green curve) at an AUROC of 0.97. Recurrent top 4 genes from our progressive transcriptome Lasso are visualized as a heatmap in **Figure 5F** clearly capturing the incremental gradient in gene expression between controls, ET, PV and MF. These include *ADAMTS3* (ADAM metallopeptidase protease(Mead and Apte, 2018) with likely roles in *VEGF* signaling, tissue remodeling and expression of related collagens through profibrotic *PAR1* and *TGF-β* signaling), *PSMB5* (implicated in proteasomal degradation/UPR activation(Wang et al., 2017) and identified previously in MPNs (Skov et al., 2010)), *NT5C* related to *PI3K* signaling (Fruman et al., 2017; Moniz et al., 2017) and *SUPT6H/SPT6(Bres et al., 2008; Frydman et al., 2020)*, a tumor-initiating histone chaperone associated with chromatin remodeling. Lasso-selected candidate markers capture the underlying MPN pathology and offer potential therapeutic targets.

A key aspect of the layered Lasso modeling demonstrated here is our use of an approach that can be developed in future work to incorporate additional predictors to MF (e.g. *JAK2* V617F allele burden (Vannucchi et al., 2008) or other genetic variants (Tefferi et al., 2016) beyond the driver mutations). **Figure S5** demonstrates this through a second base model for prediction to MF from ET or PV alone (no transcriptome, but platelet count and hemoglobin levels as two additional clinical parameters that were available for all patients in addition to age, gender and driver mutation status).

## Discussion

Here, we present a comprehensive catalog of the platelet transcriptome in chronic progressive MPN with immediate relevance to defining subtype-specific molecular differences and predicting the advanced phenotype, myelofibrosis. Recent data(Williams et al., 2020) identify the timing of MPN driver mutation acquisition to be very early in life, even before birth, with life-long clonal expansion and evolution. These new findings highlight the importance of early progression biomarkers and the substantial opportunity for early detection and intervention strategies in these disorders.

Our analyses confirm and extend many important observations made previously either in vitro (LaFave and Levine, 2016; Merlinsky et al., 2019; Osorio et al., 2016), in other transcriptome or microarray analyses (Gangaraju et al., 2020; Guo et al., 2019; Rampal et al., 2014; Rontauroli et al., 2021; Skov et al., 2012; Wong et al., 2019) including our own early work(Krishnan et al., 2017), or using animal models (Dunbar et al., 2017; Matsuura et al., 2020; Mullally et al., 2013; Wen et al., 2015). By highlighting intersecting mechanisms in transcription across MPN, and by annotating MPN subtype-specific gene signatures, this dataset facilitates predictive machine learning algorithms, that aid in MPN classification and potential prognostication.

The platelet transcriptome is significantly reprogrammed in the MPN setting, with a wealth of transcript associations that may be missed in using conventional tissue sources such as serum, plasma, whole blood or bulk bone marrow. While previous bulk RNA-seq studies on MPNs by us and others analyzed far fewer samples (Guo et al., 2019; Krishnan et al., 2017; Skov et al., 2011; Skov et al., 2012), select MPN subtypes (Gangaraju et al., 2020; Guo et al., 2019), or non-specific source tissue (Schischlik et al., 2019; Skov et al., 2011; Skov et al., 2012) that may be underpowered (Cummings et al., 2017) for candidate genes, we here analyzed, by next generation RNA sequencing, 120 purified platelet samples from healthy controls and all three subtypes; and identified clinically interpretable transcriptomic signatures for each of the three subtypes. Each subtype showed both overlapping and progressively divergent transcriptional pathways, suggesting both a shared signature across all MPN, and unique biological trajectories. Pathway-enrichment analyses confirmed the existence of a shared inflammatory milieu (Barbui et al., 2011; Geyer et al., 2015; Hasselbalch and Bjorn, 2015; Koschmieder and Chatain, 2020; Marin Oyarzún and Heller, 2019; Mughal et al., 2019; Skov et al., 2011) among MPN. We also confirmed that the JAK1/JAK2 inhibitor ruxolitinib was associated with inhibition of inflammatory as well as interferon-mediated signaling pathways. Additional previously undescribed insights into the mechanisms of action of RUX in MF included genes implicated in protein maturation, chaperone-mediated protein complex assembly, and circadian rhythm. These and other gene signatures and pathways identified may help guide candidate drugs to be used alone or in combination with RUX for the treatment of MPNs. Whether MPN oncogenic driver mutations increase inflammation or mutations are acquired in response to inflammatory stimuli is unclear from this work and remains an active area of investigation (Curto-Garcia et al., 2020; Hasselbalch and Bjorn, 2015; Koschmieder and Chatain, 2020; Lussana et al., 2017).

The 10 genes most significant (FDR < 0.001) of the commonly expressed genes across MPN indicated a gradation in platelet gene expression, with overlapping signatures in ET and PV (*e.g. IFITM2, MYBL2)* and a substantial difference with MF (*e.g. CREB3L1, CALR*) that was independent of driver mutation status or treatment. Hence, while over 1500 genes were commonly differentially expressed across MPN, their abundance and function could differ between subtypes. The nature of the separation of transcriptomic clusters between ET, PV and MF suggest also that they represent diverse cell states along a continuous spectrum of MPN, in line with the clinical overlap of these neoplasms.

Another observation relates to the association of the differential genes with signaling pathways: as indicated above, all three MPN subtypes showed a positive enrichment in immune modulation pathways, independent of mutational status. Whether this response reflects a causal effect of inflammation on bone marrow biology remains to be elucidated. Indeed, the platelet transcriptomic signatures could also reflect inter-cell interactions of platelets with other immune cells, including as transient aggregates with neutrophils, granulocytes and dendritic cells. Nevertheless, observations that MPN transcriptomic biomarkers correlated robustly with immune factors such as type I/II interferons and dysregulation of interleukin-dependent inflammatory responses across ET, PV and MF suggest opportunities for use of these and other subtype-specific genes as biomarkers for prognosis as well as design of therapies and prediction of response.

Our data closely overlaps with recent MPN platelet studies: thrombo-inflammatory signatures in PV from Gangaraju and Prchal et al (Gangaraju et al., 2020) (*BCL2, CXCL1, MMP7, PGLYRP1, CKB, BSG, CFL1* and more) or fibrosis-associated signatures in MF from Guo et al. (*CCND1, H2AFX, CEP55* and several others collectively reflected in the external validation of our Lasso algorithm).

Most notably in our data on MF, high expression of ER stress and unfolded protein response (UPR) biomarkers *(e.g. CREB3L1, CALR)* associated with impaired proteostasis signaling; and emerged as a key feature of MPN pathobiology. Indeed, recently published work from distinct research groups (Liu et al., 2020; Osorio et al., 2016),(LaFave and Levine, 2016) highlight protein quality control in ER-associated degradation and proteostasis (Osorio et al., 2016),(LaFave and Levine, 2016) deregulation as a primary effector of myeloid transformation highlighting the importance of protein homeostasis for normal hematopoiesis. These findings too are in line with reports (Kaushik and Cuervo, 2015) implicating chronic ER stress, malfunctioning protein quality control, and loss of proteostasis as aggravating factors in age-related disorders.

Most importantly, platelet gene expression profiling in MPN offers directions for prediction of myelofibrosis. Applying machine-learning algorithms of LASSO penalized regression under two conditions of external validation (Moons et al., 2012): temporal (using our two cohort design) and geographical (independently published datasets on healthy donors(Guo et al., 2020; Rondina et al., 2020) and MF (Guo et al., 2020), we uniquely discriminate MPN subtypes from each other, and healthy controls using three model types and predict MF at high accuracy. The highest performing model used a set of progressively differentiated MPN genes at an area under the (ROC) curve of 0.96 (temporal) and 0.97 (geographical); and rendered a core signature of <5 candidate markers as top predictors of disease progression. It will be of interest to determine what machine-learning algorithms based on a defined platelet gene expression classifier on potential new MPN datasets (ideally longitudinal) can be used to more precisely predict the probability and/or timing of an individual’s risk of progression from ET/PV to secondary MF.

In conclusion, using platelet transcriptome profiling, we observed dynamic shifts in MPN immune inflammatory profile and preferential expression of interferon-, proliferation-, and proteostasis-associated genes as a progressive gradient across the three MPN subtypes. Our findings highlight that MPN progression may be influenced by defects in protein homeostasis (impaired protein folding and an accumulation of misfolded proteins within the endoplasmic reticulum) and an abnormal integrated stress response – consistent with recent studies^57,^(Liu et al., 2020; Osorio et al., 2016),(LaFave and Levine, 2016) indicating dysregulated proteostasis as a primary effector of myeloid transformation. While this particular work has been focused on the overarching progressive platelet transcriptome across MPNs, these data open an important avenue for utilizing platelet RNA signatures to better understand specific MPN complications such as the risk of thrombosis and bleeding, or fibrosis and transformation to AML. Altogether, this study demonstrated in chronic MPNs provides a comprehensive framework for exploiting the platelet transcriptome and may inform future studies toward mechanistic understanding and therapeutic development in MPNs, and potentially other age-related disorders.

### Limitations of the study

There are several limitations to our study. First, our data are not longitudinal by design but rather the closest practical alternative of cross-sectional snapshots of all three MPN subtypes with the goal of achieving a well-powered dataset in these chronic disorders. In this regard, the progressive or progression terminology used here refer strictly to trends in gene expression and do not imply study of longitudinal clinical progression. Therefore, subsidiary longitudinal evaluation of the disease as well as treatment markers identified here is warranted. Second, our focus for this study has been on the platelet transcriptome alone. Future investigations focused on ascertaining overlap between our platelet-derived molecular alterations with those of other cell types, specifically, parent megakaryocytes, CD34+ cells, granulocytes/immune cells and even whole blood will be required to identify additional functional aspects of bone marrow pathology. Such integrative analyses may also necessitate advanced systems genomics methods that compare or combine data without biases and batch effects inherent in each cell data type. Third, we recognize that our choice of ribosomal RNA depletion to home in on platelet mRNA signatures leaves out additional diversity in the platelet RNA repertoire (and will be important future work). Lastly, in our Lasso predictive modeling, we demonstrate two rigorous approaches of external validation (temporal and geographical) and identify a core signature toward MPN risk stratification or early detection of progression. Yet, substantive future biological and computational validations are needed in order to advance our findings toward clinical decision making or personalized medicine.

## Methods

### Ethical Approval

All MPN peripheral blood samples were obtained under written informed patient consent and were fully anonymized. Study approval was provided by the Stanford University Institutional Review Board. All relevant ethical regulations were followed.

### Subjects and Specimen Collection

We collected blood from ninety-five MPN patients enrolled in the Stanford University and Stanford Cancer Institute Hematology Tissue Bank from December 2016-December 2019 after written informed consent from patients or their legally authorized representatives (Stanford IRB approval #18329). Eligibility criteria included age ≥18 years and Stanford MPN clinic diagnosis of essential thrombocythemia, polycythemia vera or myelofibrosis (defined using the consensus criteria at the time of this study). We use the term ‘myelofibrosis’ to encompass both primary myelofibrosis and myelofibrosis that evolved from essential thrombocythemia or polycythemia vera. Electronic medical records review of all subjects was performed by the clinical consultants (J.G. and L.F.), study data manager (C.P.), and the study principal investigator (A.K.). For controls, blood was collected from twenty-one asymptomatic adult donors selected at random from the Stanford Blood Center. All donors were asked for consent for genetic research. For both MPN patients and healthy controls, blood was collected into acid citrate-dextrose (ACD, 3.2%) sterile yellow-top tubes (Becton, Dickinson and Co.) and platelets were isolated by established(Amisten, 2012; Campbell et al., 2018; Middleton et al., 2019; Rowley et al., 2011) purification protocols. Blood was processed within 4h of collection for all samples. The time from whole blood collection to platelet isolation was similar between healthy donors and MPN patients.

### Platelet Isolation

Human platelets were isolated and leuko-depleted using established methods ((Amisten, 2012; Campbell et al., 2018; Rowley et al., 2011) with excellent reproducibility(Campbell et al., 2018; Davizon-Castillo et al., 2020; Manne et al., 2020; Middleton et al., 2019; Rondina et al., 2013) resulting in a highly purified population of fewer than 3 leukocytes/10^7^ platelets (>99.9% purity) as counted by hemocytometer. Briefly, the ACD-tube whole blood was first centrifuged at 200xg for 20min at room temperature (RT). The platelet-rich plasma (PRP) was removed and Prostaglandin E1 was added to the PRP to prevent exogenous platelet activation. The PRP was then centrifuged at 1000xg for 20min at RT. The platelet pellet was re-suspended in warmed (37 deg C) PIPES saline glucose (PSG). Leukocytes were depleted using CD45+ magnetic beads (Miltenyi Biotec). Isolated platelets were lysed in Trizol for RNA extraction. Post RNA-seq analysis of an index leukocyte transcript (*PTPRC; CD45*) confirmed that the samples were highly purified platelet preparations (subsequent bioinformatic analyses also adjusted for PTPRC expression for absolute removal of any CD45 expression in our analyses). Two reference markers of platelet activation, *P-selectin (SELP)* and *Glycoprotein IIbIIIA (CD41/ITGA2B)* were expectedly higher in all MPN than healthy controls but were not statistically significantly different between MPN subtypes; indicating that any expression difference was not due to experimental artefacts. In addition, we know from prior work (J.R.) that regardless of activation status, RNA-seq reliably estimates mRNA expression patterns in platelets. We also know from rigorous prior work(Gnatenko et al., 2003; Raghavachari et al., 2007; Weyrich and Zimmerman, 2003) that several abundant platelet mRNAs are well-known leukocyte or red cell transcripts; and do not immediately imply contamination by these classes but rather that platelets express gene products that are also present in other cell lineages. Sixty-five of our top 100 abundant platelet transcripts matched exactly with those of the top 100 abundant genes from the three previous studies cited(Gnatenko et al., 2003; Rowley et al., 2011); and a composite pathway analysis with the top 100 abundant genes from this as well as the previous studies matched identically.

### Next Generation RNA Sequencing

For next generation RNA-sequencing (RNA-seq), 1x10^9^ isolated platelets were lysed in Trizol and then DNAse treated. Total RNA was isolated, and an Agilent bio-analyzer was used to quantify the amount and quality. The RNA yield was estimated by measuring absorbance at 260 nm on the Nanodrop 2000 (Thermo Fisher), and RNA purity was determined by calculating 260/280 nm and 260/230 nm absorbance ratios. RNA integrity was assessed on the Agilent Bioanalyzer using the RNA 6000 Nano Chip kit (Agilent Technologies). An RNA integrity number (RIN) was assigned to each sample by the accompanying Bioanalyzer Expert 2100 software. To control for variable RNA quality, RNA sequencing was only performed for samples with a RIN score of 7 or higher. RNA-seq libraries were constructed with removal of ribosomal RNA using the KAPA Stranded RNA-Seq kit with RiboErase (Roche). The RNA extraction and library preparation were performed by the same technician to minimize confounding effects. cDNA libraries were constructed following the Illumina TrueSeq Stranded mRNA Sample Prep Kit protocol and dual indexed. The average size and quality of each cDNA library were determined by the Agilent Bioanalyzer 2100, and concentrations were determined by Qubit for proper dilutions and balancing across samples. Twelve pooled samples with individual indices were run on an Illumina HiSeq 4000 (Patterned flow cell with Hiseq4000 SBS v3 chemistry) as 2 X 75bp paired end sequencing with a coverage goal of 40M reads/sample. Output BCL files were FASTQ-converted and demultiplexed.

### Platelet Transcriptome Analysis

Picard, Samtools, and other metrics were used to evaluate data quality. Processed reads were aligned against the reference human transcriptome GRCh37/hg19 using RSEM(Li and Dewey, 2011) and bowtie2(Langmead and Salzberg, 2012), and expression at gene level determined by calculating raw gene count. Only genes that passed expression threshold were used; genes were considered expressed if, in all samples, they had at least 10 counts (genes with low counts are automatically filtered by built-in functions in DeSeq2, see below). A total of 12,924 genes were considered expressed. Gene expression data was library-size-corrected, variance-stabilized, and log2-transformed using the R package DESeq2(Love et al., 2014). We refer to this version of the data as “raw data” as it is not corrected for any confounders of gene expression variability. DESeq2 was used to call differential expression while adjusting for patient age, gender and treatment as confounding variables and controlling for multiple comparisons using the Benjamini-Hochberg defined false discovery rate (FDR). Significant variance in expressed transcripts were pre-specified as those transcripts with an FDR <0.05 and a log2 fold change ≥ 0.5 in MPN, as compared to healthy controls (the entire differential transcriptome was applied in the instances of downstream Gene Set Enrichment Analysis and the Lasso prediction modeling).

### Statistical analysis

Continuous data were summarized as medians and IQRs and categorical data are presented as frequencies and percentages. To compare differences in clinical variables between healthy controls and MPN subtypes (ET, PV and MF), we use box and whisker plots and conduct pairwise Wilcoxon signed ranked tests. For unsupervised clustering and visualization, we performed principal component analyses (identifying MPN subtypes by color, treatment by filled or open circles, and *JAK2* mutation status by shape) using built-in functions of the DeSeq2 R package. We generated a heatmap of all of the top highly significant genes (FDR < 0.01) using the pheatmap R package and its built-in functions for hierarchical cluster analysis on the sample-to-sample Euclidean distance matrix of the expression data. All analyses were performed using the R studio interface.

### Pathway/Gene set enrichment analysis for differentially expressed (DE) genes

Gene set enrichment analysis (GSEA)(Subramanian et al., 2005) was performed on the entire DE gene set for each MPN subtype, using the Cancer Hallmarks gene sets from MSigDB(Liberzon et al., 2015). The ‘GSEA Pre-ranked’ function was used with a metric score that combines fold change and adjusted p-value together for improved gene ranking. We used default settings with 10,000 gene set permutations to generate *p* and *q* values, and compared MPN subtypes in the overall cohort, and the ruxolitinib-treated subgroup and the ruxolitinib-naive subgroup separately. In these analyses, to allow for a broad comparison, we assessed all transcripts that were differentially expressed according to FDR/adjusted p < 0.25 as recommended by the authors of GSEA (Subramanian et al., 2005).

### Predictive model generation and external validation

At the conception of this study late 2016, and our early work(Krishnan et al., 2017), we did not identify any publicly available RNA sequencing data on MPN platelets. This prompted our specific two-cohort design for the express purpose of temporal external validation as an essential step in rigorous prediction modeling(Moons et al., 2012). A subsequent independent publication(Guo et al., 2020) facilitated an additional geographically independent external validation of our model.

We used Lasso penalized regression(Tibshirani, 1996) for our model to predict MF from the either healthy controls, ET or PV. Among a variety of statistical machine learning algorithms that have been used in prediction modeling, Lasso is favored for its flexibility and simplicity; and its ability to identify the least set of significant factors from high dimensional data. We evaluated platelet transcriptomic features with clinical features (age, gender and mutation status for the entire dataset including healthy donors, and in MPN patients alone, platelet and hemoglobin values). Normalized gene counts data were split into training (used for constructing multinomial logistic models) and validation (used for model evaluation and generalization) cohorts. Separately, we assessed the progressive and monotonic upward or downward trend in gene expression, we applied the Mann-Kendall trend test (multiple comparison adjusted with the Benjamini-Hochberg method) to normalized gene counts and identified statistically significant progressive genes across all three MPN subtypes.

Three multinomial logistic models were constructed: first, with Lasso selected predictors from all genes, second, with Lasso selected predictors from progressive genes and third, a baseline model using age, gender and mutation status (*JAK2 and CALR*) as predictors. Model outputs correspond to probabilities of having a CTRL, ET, PV or MF phenotype (sum of these four probability values totaling 1). Potential interpretation of these probabilities includes MPN risk assessment, *e.g.* a patient with higher probabilities of PV and MF would indicate higher risk than one with higher probabilities of CTRL or ET. The (Guo et al., 2020) dataset on MF platelet RNA seq served as an independent test set (Fig. 5D schematic) while data from our cohorts at Stanford and additional external data on healthy donors from (Rondina et al., 2020) constituted an integrated training cohort (R package Limma was applied for bioinformatic correction of any batch effects).

ROC curves were used to evaluate the different prediction models and discriminate outcomes. ROC curves demonstrate the trade-off between true positive and false positive rates, ideal being high true positive rate (sensitivity) and low false positive rate (specificity) the area under the curve (AUROC) as close to 1 as possible. True positive rate (TPR) is defined as correctly predicting an MF patient as MF; and false positive rate (FPR) as falsely predicting a non-MF patient as MF.

## Authorship Contributions

A. Krishnan, J. Gotlib and J. Zehnder conceived of the overall study. A. Krishnan designed the study and secured funding that initiated this research. L.F., C.P., and J.G. provided samples and clinical annotation and reviewed the clinical data. A.K. designed the experimental plan with input from J.R., H.M., J.G. and J.Z. A.K. coordinated, performed and oversaw the sample acquisition and processing. V.N. performed RNA isolation and library preparation. A.K. coordinated and oversaw sample sequencing. Z.S., W.D. and A.K. performed and interpreted the computational analyses. A.K. J.R., H.M., J.G. and J.Z. interpreted the data. A.K. wrote and edited the manuscript; C.P., W.D., J.R., H.M., J.G. and J.Z. critically reviewed and edited the manuscript. ^+^J.G. and J.Z. contributed equally. All authors approved the final manuscript.

## Acknowledgements

This work was funded by US National Institutes of Health grants 1K08HG010061-01A1 and 3UL1TR001085-04S1 (research re-entry award) to A.K and the Charles and Ann Johnson Foundation to J.G. A.K. would like to thank mentors, Profs. Richard Becker (University of Cincinnati, OH), Andrew Weyrich (University of Utah, UT), Harry Greenberg (Stanford University, CA), Rob Tibshirani (Stanford University, CA) and Stephen Montgomery (Stanford University, CA) for their support in her unique return to research and NIH funding after a 5-year hiatus that then led to the work outlined in this manuscript. All authors thank the patients at the Stanford Cancer Center for their generous participation in this research, and the Stanford Functional Genomics Facility for genomic data storage. Authors also thank Drs. Belinda Guo and Wendy Erber (University of Western Australia, Australia) for the gene counts file of their published data (PMID 31426129) used for our independent validation. A.K. extends special thanks to colleagues Dr. Kellie Machlus (Harvard University, MA), and Dr. Eric Pietras (University of Colorado, CO) for their critical reading of the manuscript.

## Data Sharing Statement

RNA-sequencing data from this work (original FASQ files from paired-end sequencing of all 120 samples) will be deposited to the NIH genomic data repository dbGAP under public accession # PHS-0021-21. v1.P1. Previously published RNA-sequencing data used in this work as geographically independent validation cohorts are from Rondina et al (PMID 31852401, healthy donors) and Guo et al (PMID 31426129, MF patients and healthy donors). Source data from the work of Rondina et al is publicly available at NIH NCBI Bioproject ID 531691; and that of Guo et al is secured through reaching the corresponding author, Dr. Wendy Erber.

A private link for editors and reviewers will be made available at a Stanford web-based data repository.

## Ethics Declarations/Competing Interests

The authors declare no competing interests.

## Supplementary Figure Legends

**Figure S1:**
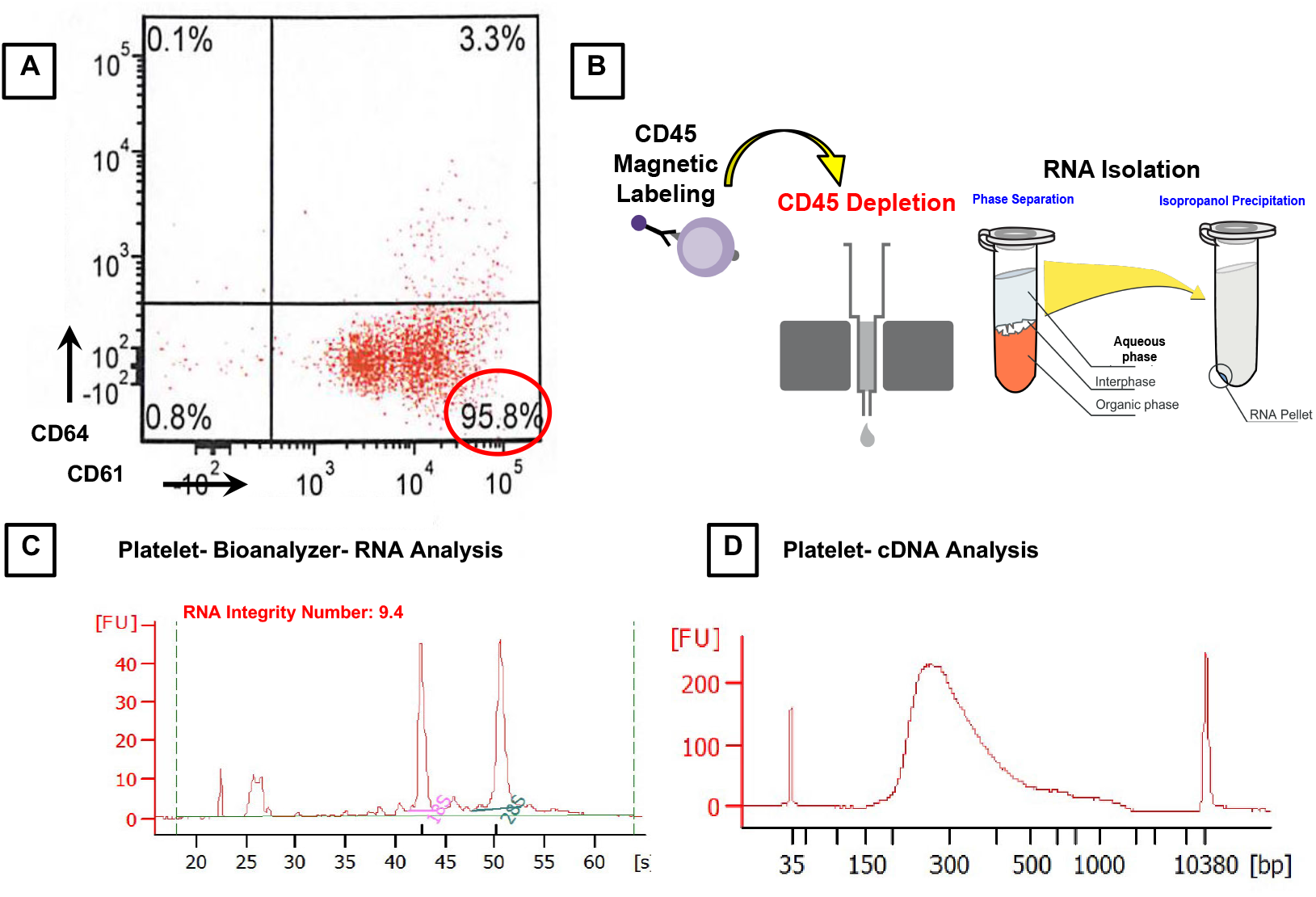
Platelets isolated for RNA-sequencing. **A**. Flow cytometry analysis for purity of platelets (CD61+) isolated from whole blood. Population (quadrant 3, outlined in red) showing 96% pure platelets prior to enrichment by depletion of CD45 leukocytes. **B.** CD45 magnetic microbeads were used to deplete leukocytes, further enriching platelet population. Platelets enriched from magnetic sorting were confirmed for ∼99% platelet purity using a Cell-Dyn Hematology Analyzer prior to isolation of RNA. Standard trizol RNA extraction protocol was used to extract RNA. **C.** Representative electropherogram results from Agilent Bioanalyzer showing RNA isolated from platelets with an RNA Integrity Number of 9.4**. D**. Electropherogram showing a clean product following cDNA synthesis and amplification. Similar procedure was carried out for all samples and by the same individual.

**Figure S2:**
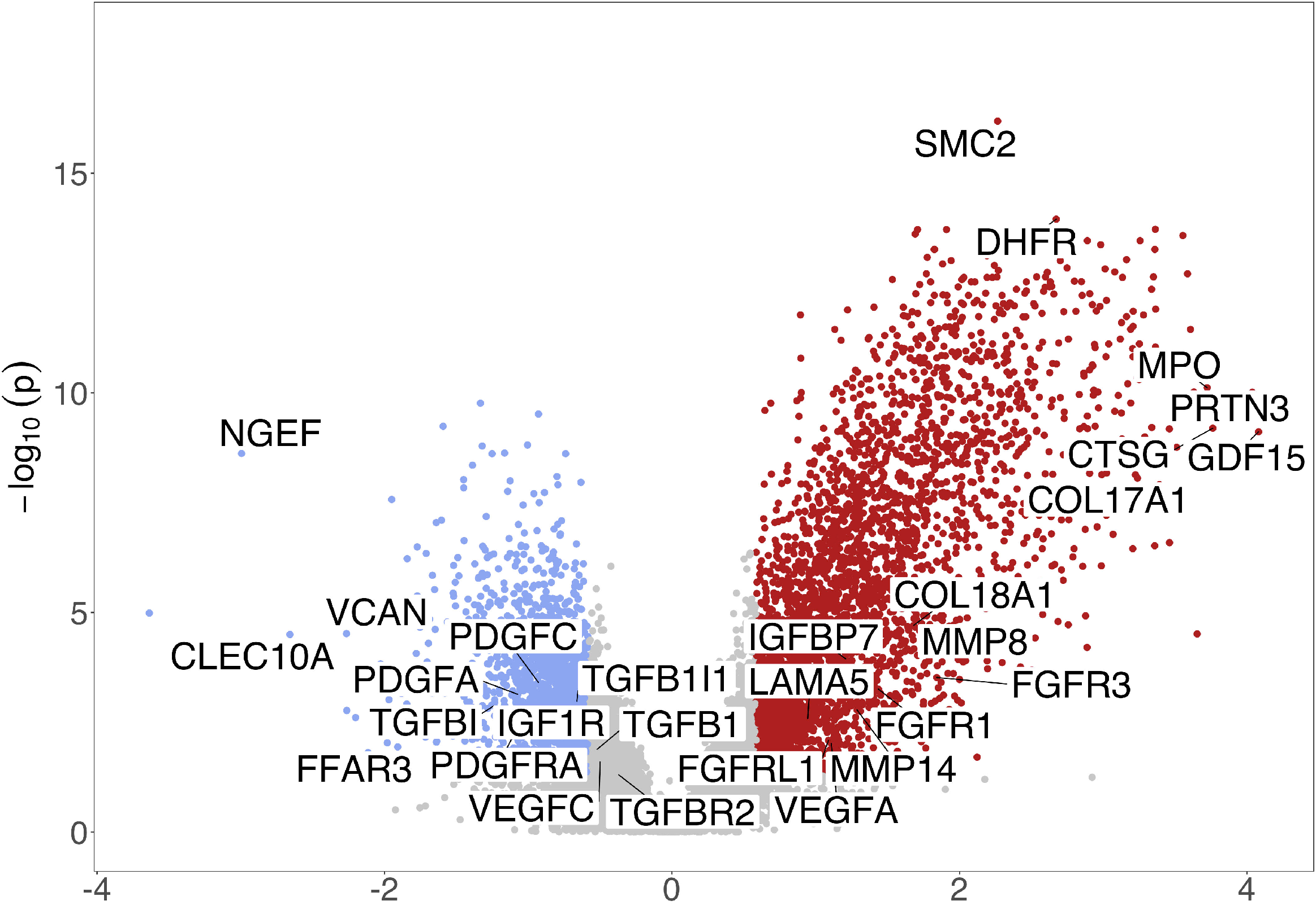
Volcano plots (showing statistical significance, negative log10 of p-values, versus log2 fold change of each gene) of differential gene expression analysis of a sub-cohort of MF versus ET and PV patients combined. Significant up- and down-regulated genes are those with p-values (FDR) smaller or equal to 0.05 and absolute value of fold changes larger or equal to 1.5. Possible mediators of fibrosis-associated genes are highlighted and include pro-fibrotic growth factors and other matrix proteins.

**Figure S3:**
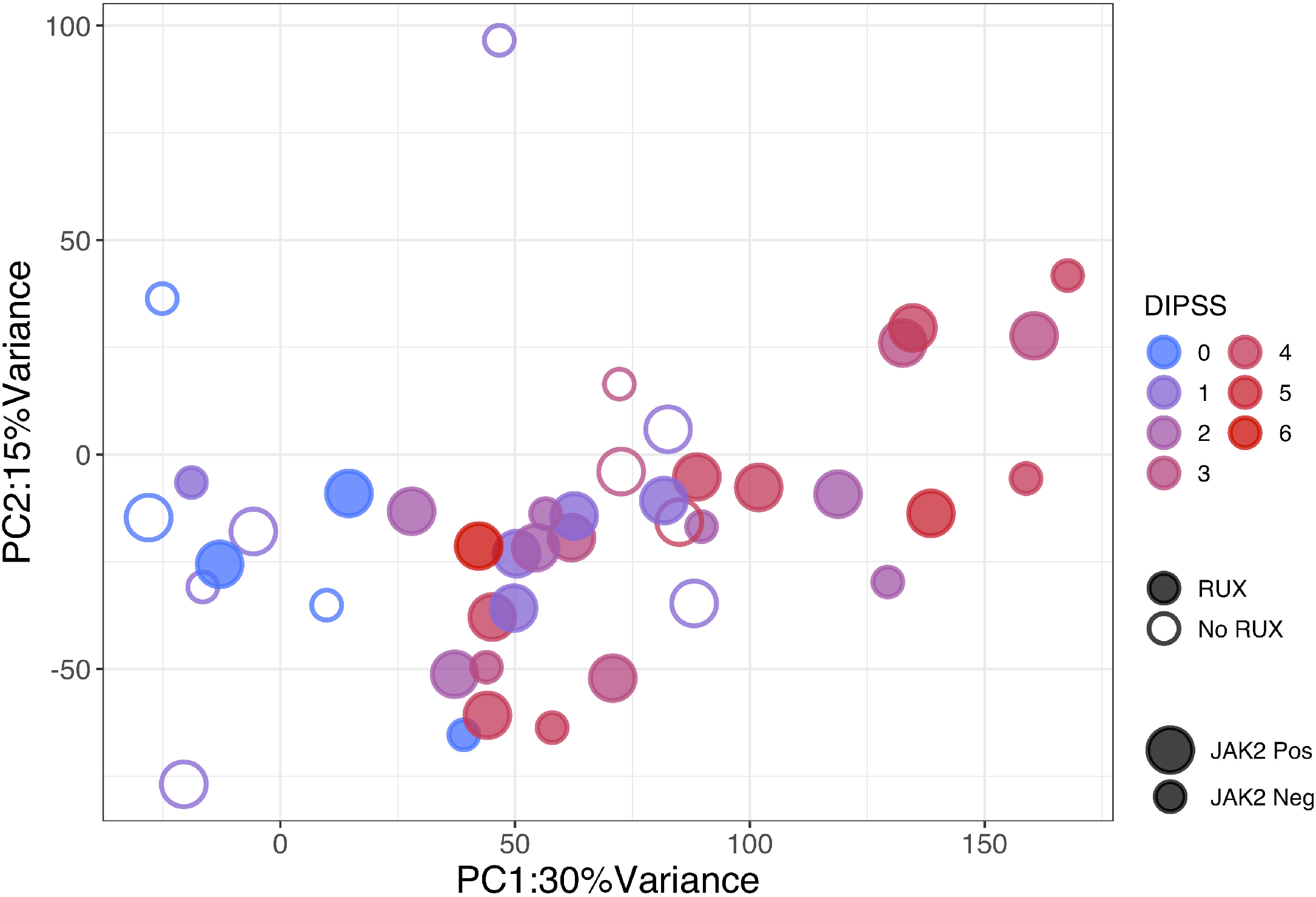
Correlation of MF platelet transcriptome with disease risk by Dynamic International Prognostic Scoring System (DIPSS) **A**, Unsupervised principal component analysis (PCA) of MF (n=42) normalized platelet gene expression counts adjusted for age, gender, treatment and experimental batch. PC1 and PC2 colored by DIPSS score (0-6 denoting increasing MF risk). Circles filled or open mark presence or absence of ruxolitinib treatment; and size of circles, smaller or larger, indicate presence or absence of *JAK2* mutation.

**Figure S4:**
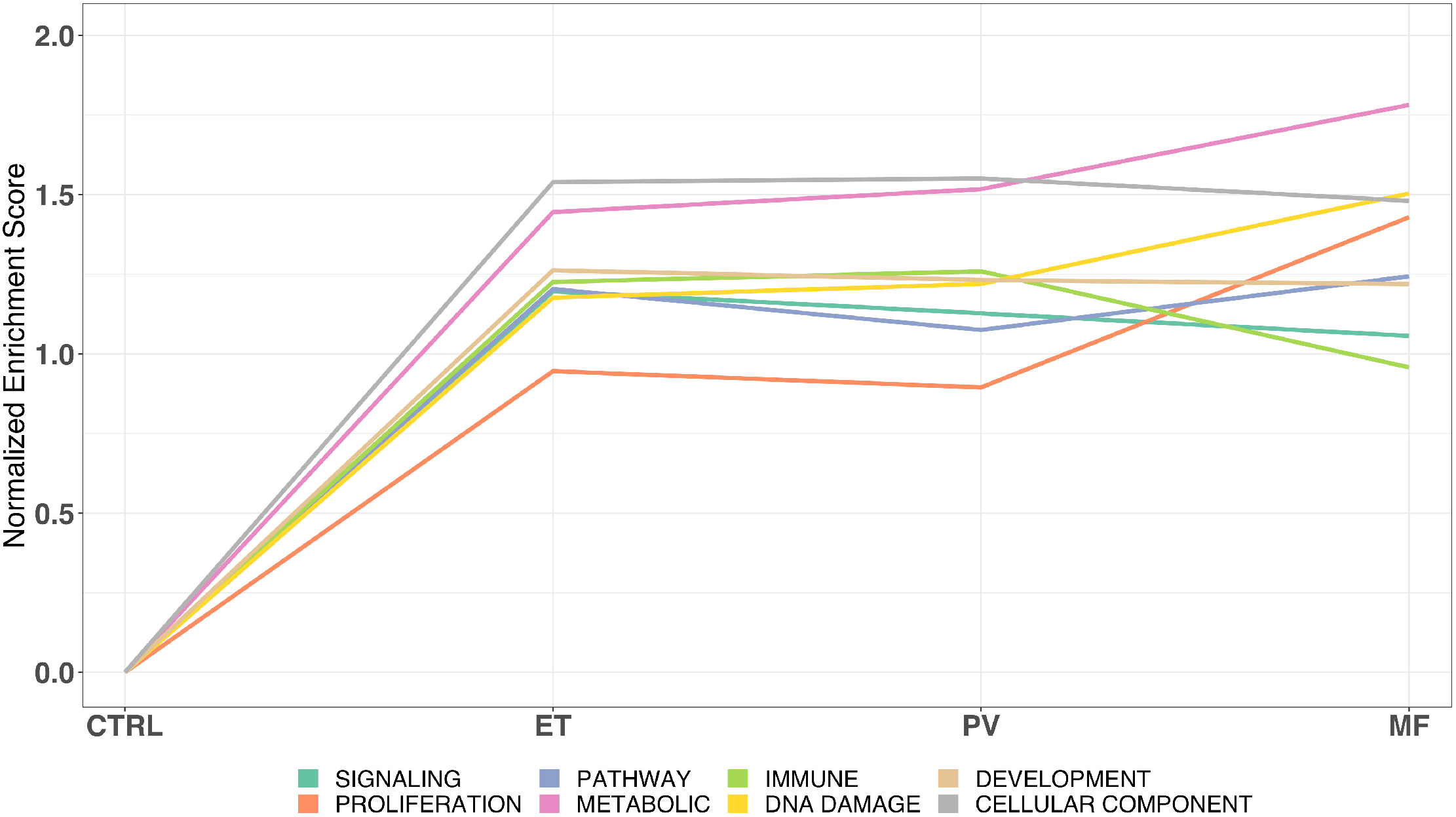
Relative molecular trajectories of MPN progression. The relative enrichment of MPN molecular pathways reflecting MPN progression within specific biological process categories.(Liberzon et al., 2015) Each color-indicated line represents a pathway category. *x*-axis captures sample type from controls (CTRL) to ET, PV and MF, and the y-axis gives the normalized enrichment score, which reflects the degree to which each pathway is over or under-represented at the top or bottom of the ranked list of differentially expressed genes, normalized to account for differences in gene set size and in correlations between gene sets and the expression data set.

**Figure S5:**
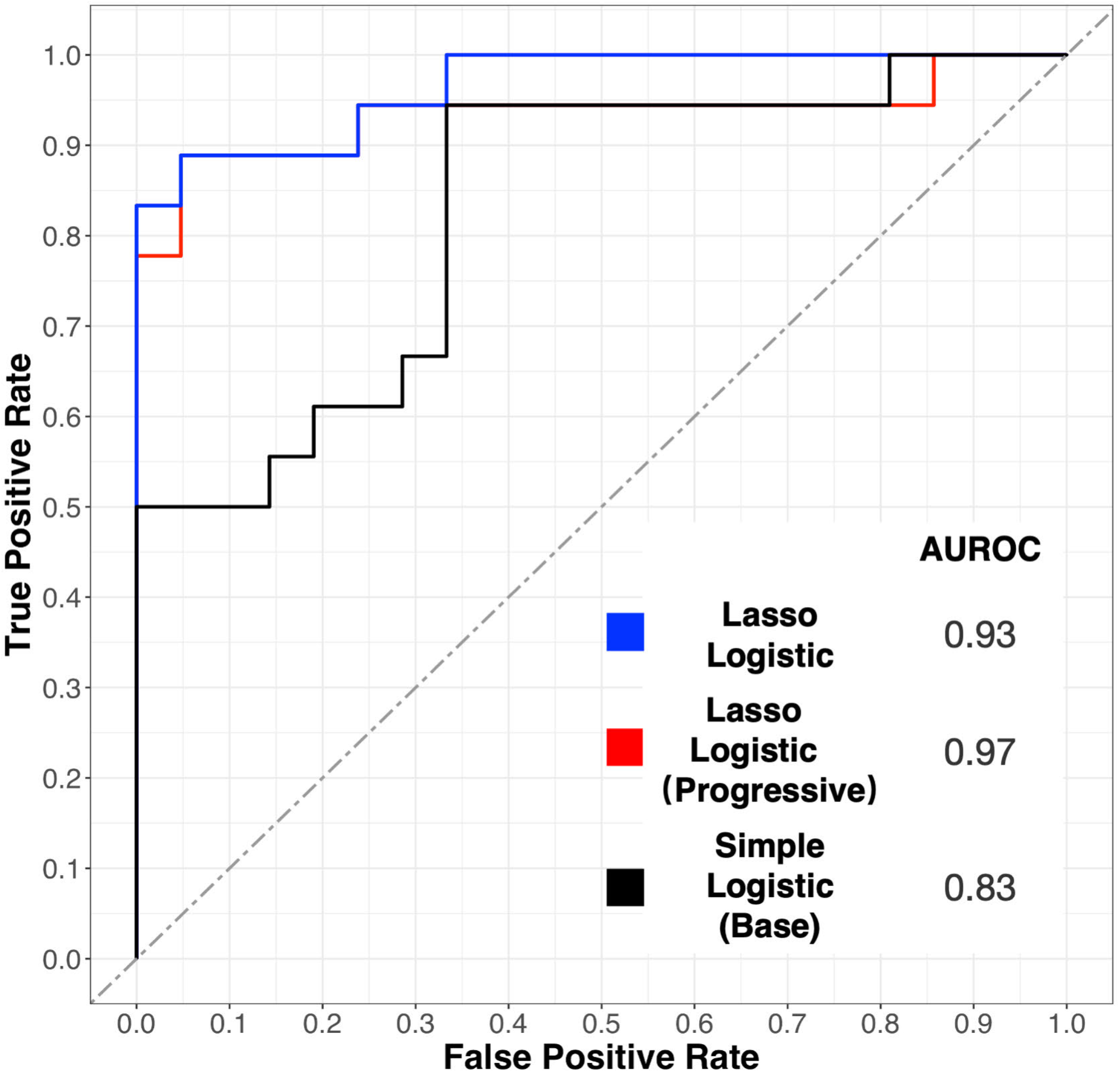
Receiver Operating Curves (ROC) toward MF prediction from ET or PV alone with temporal validation and three layered models: i) baseline, with no gene expression data but patient age, gender and mutation status (as Figure 5E) and an additional two clinical variables: platelet count and hemoglobin; ii) adding entire MPN platelet transcriptome to the above baseline and iii) adding MPN progressive genes alone. Outperformance of the progressive transcriptome model (red curve, AUROC=0.97) vis-a-vis the entire transcriptome dataset (blue curve, AUROC=0.93) and the baseline model without gene expression (black curve, AUROC=0.83).

**Figure S6:**
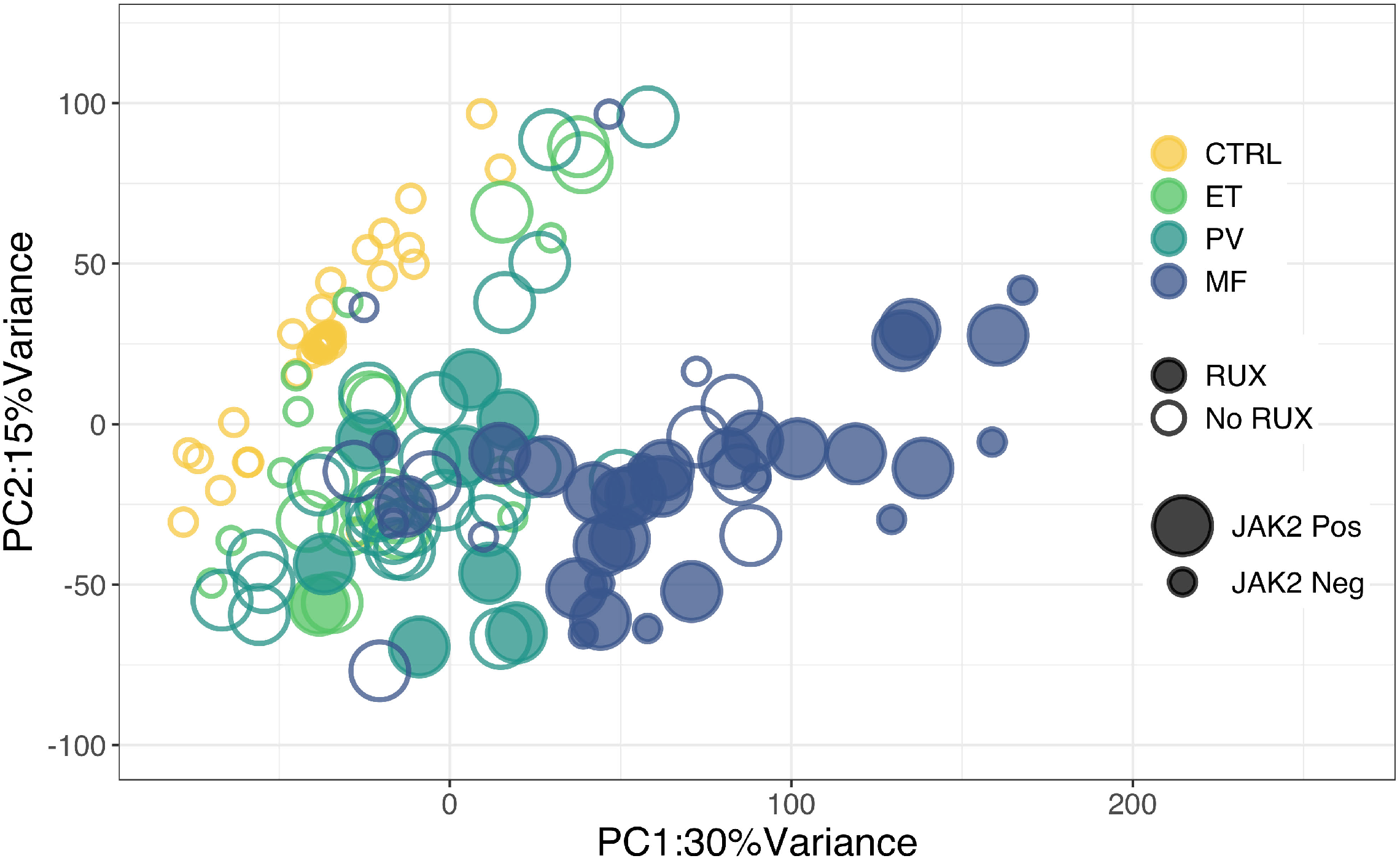
Unsupervised principal component analysis of platelet transcriptomic data from Stanford MPN & healthy donor cohorts (n=120) integrated with that of independently published healthy donors (n=31) (Rondina et al., 2020). Colors indicate controls (n=52, yellow): ET (n=24, top, light green), PV (n=33, middle, dark green) and MF (n=42, bottom, dark blue). Circles filled or open mark presence or absence of ruxolitinib treatment; and size of circles, smaller or larger, indicate presence or absence of *JAK2* mutation.

## Table Descriptions

**Table S1A.**
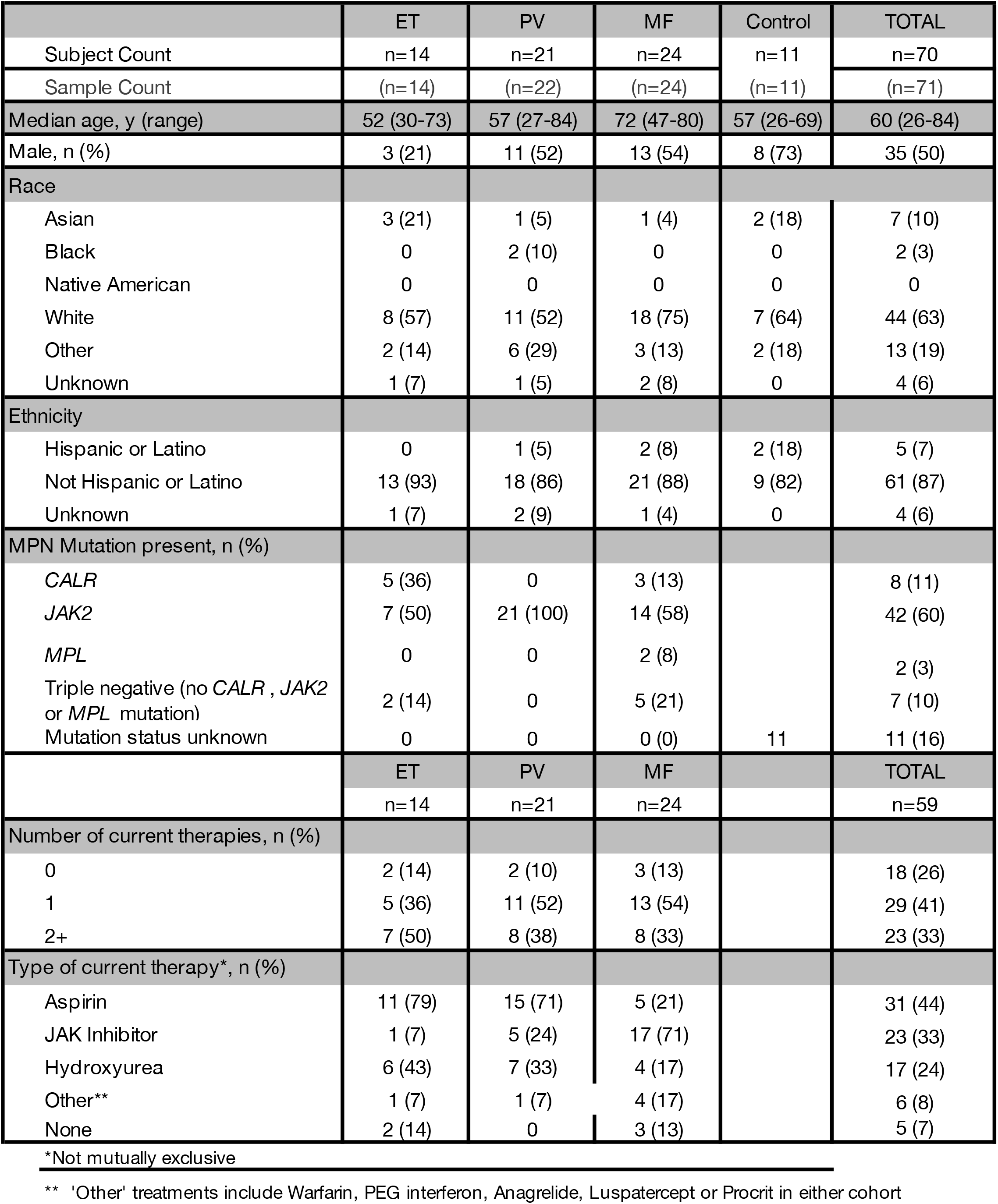
Characteristics Cohort1.

|  | ET | PV | MF | Control | TOTAL |
| --- | --- | --- | --- | --- | --- |
| Subject Count | n=14 | n=21 | n=24 | n=11 | n=70 |
| Sample Count | (n=14) | (n=22) | (n=24) | (n=11) | (n=71) |
| Median age, y (range) | 52 (30-73) | 57 (27-84) | 72 (47-80) | 57 (26-69) | 60 (26-84) |
| Male, n (%) | 3 (21) | 11 (52) | 13 (54) | 8 (73) | 35 (50) |
| Race |  |  |  |  |  |
| Asian | 3 (21) | 1 (5) | 1 (4) | 2 (18) | 7 (10) |
| Black | 0 | 2 (10) | 0 | 0 | 2 (3) |
| Native American | 0 | 0 | 0 | 0 | 0 |
| White | 8 (57) | 11 (52) | 18 (75) | 7 (64) | 44 (63) |
| Other | 2 (14) | 6 (29) | 3 (13) | 2 (18) | 13 (19) |
| Unknown | 1 (7) | 1 (5) | 2 (8) | 0 | 4 (6) |
| Ethnicity |  |  |  |  |  |
| Hispanic or Latino | 0 | 1 (5) | 2 (8) | 2 (18) | 5 (7) |
| Not Hispanic or Latino | 13 (93) | 18 (86) | 21 (88) | 9 (82) | 61 (87) |
| Unknown | 1 (7) | 2 (9) | 1 (4) | 0 | 4 (6) |
| MPN Mutation present, n (%) |  |  |  |  |  |
| *CALR* | 5 (36) | 0 | 3 (13) |  | 8 (11) |
| *JAK2* | 7 (50) | 21 (100) | 14 (58) |  | 42 (60) |
| *MPL* | 0 | 0 | 2 (8) |  | 2 (3) |
| Triple negative (no *CALR* , *JAK2* or *MPL* mutation) | 2 (14) | 0 | 5 (21) |  | 7 (10) |
| Mutation status unknown | 0 | 0 | 0 (0) | 11 | 11 (16) |
|  | ET | PV | MF |  | TOTAL |
|  | n=14 | n=21 | n=24 |  | n=59 |
| Number of current therapies, n (%) |  |  |  |  |  |
| 0 | 2 (14) | 2 (10) | 3 (13) |  | 18 (26) |
| 1 | 5 (36) | 11 (52) | 13 (54) |  | 29 (41) |
| 2+ | 7 (50) | 8 (38) | 8 (33) |  | 23 (33) |
| Type of current therapy*, n (%) |  |  |  |  |  |
| Aspirin | 11 (79) | 15 (71) | 5 (21) |  | 31 (44) |
| JAK Inhibitor | 1 (7) | 5 (24) | 17 (71) |  | 23 (33) |
| Hydroxyurea | 6 (43) | 7 (33) | 4 (17) |  | 17 (24) |
| Other** | 1 (7) | 1 (7) | 4 (17) |  | 6 (8) |
| None | 2 (14) | 0 | 3 (13) |  | 5 (7) |

\*Not mutually exclusive
\*\* 'Other' treatments include Warfarin, PEG interferon, Anagrelide, Luspatercept or Procrit in either cohort

**Table S1B.** Characteristics Cohort2.

|  | ET | PV | MF | Control | TOTAL |
| --- | --- | --- | --- | --- | --- |
| Subject Count | n=10 | n=10 | n=16 | n=10 | n=46 |
| Sample Count | (n=10) | (n=11) | (n=18) | (n=10) | (n=49) |
| Median age, y (range) | 65 (31-78) | 65 (29-84) | 69 (43-83) | 57 (26-69) | 65 (26-84) |
| Male, n (%) | 5 (50) | 6 (60) | 8 (50) | 8 (80) | 27 (59) |
| Race |  |  |  |  |  |
| Asian | 1 (10) | 1 (10) | 6 (38) | 5 (50) | 13 (28) |
| Black | 0 | 0 | 1 (6) | 0 | 1 (2) |
| Native American | 0 | 0 | 0 | 2 (20) | 2 (4) |
| White | 8 (80) | 8 (80) | 7 (44) | 2 (20) | 25 (54) |
| Other | 0 | 1 (10) | 2 (13) | 1 (10) | 4 (9) |
| Unknown | 1 (10) | 0 | 0 | 0 | 1 (2) |
| Ethnicity |  |  |  |  |  |
| Hispanic or Latino | 0 | 1 (10) | 1 (6) | 1 (10) | 3 (6) |
| Not Hispanic or Latino | 9 (90) | 9 (90) | 15 (94) | 9 (90) | 42 (91) |
| Unknown | 1 (10) | 0 | 0 | 0 | 1 (2) |
| MPN Mutation present, n (%) |  |  |  |  |  |
| *CALR* | 2 (20) | 0 | 3 (19) |  | 5 (11) |
| *JAK2* | 6 (60) | 10 (100) | 11 (68) |  | 27 (59) |
| *MPL* | 2 (20) | 0 | 0 |  | 2 (4) |
| Triple negative (no *CALR*,<br>*JAK2* or *MPL* mutation) | 0 | 0 | 2 (13) |  | 2 (4) |
| Mutation status unknown | 0 | 0 | 0 | 10 | 10 (22) |
|  | ET | PV | MF |  | TOTAL |
|  | n=10 | n=10 | n=16 |  | n=36 |
| Number of current therapies, n (%) |  |  |  |  |  |
| 0 | 1 (10) | 0 | 1 (6) |  | 2 (6) |
| 1 | 7 (70) | 6 (60) | 10 (63) |  | 23 (64) |
| 2+ | 2 (20) | 4 (40) | 5 (31) |  | 11 (31) |
| Type of current therapy*, n (%) |  |  |  |  |  |
| Aspirin | 5 (50) | 7 (70) | 4 (25) |  | 16 (44) |
| JAK Inhibitor | 0 | 2 (20) | 11 (69) |  | 13 (36) |
| Hydroxyurea | 5 (50) | 4 (40) | 1 (6) |  | 10 (28) |
| Other** | 1 (10) | 2 (20) | 4 (25) |  | 7 (19) |
| None | 1 (10) | 0 | 1 (6) |  | 2 (6) |

**TableS2A:**
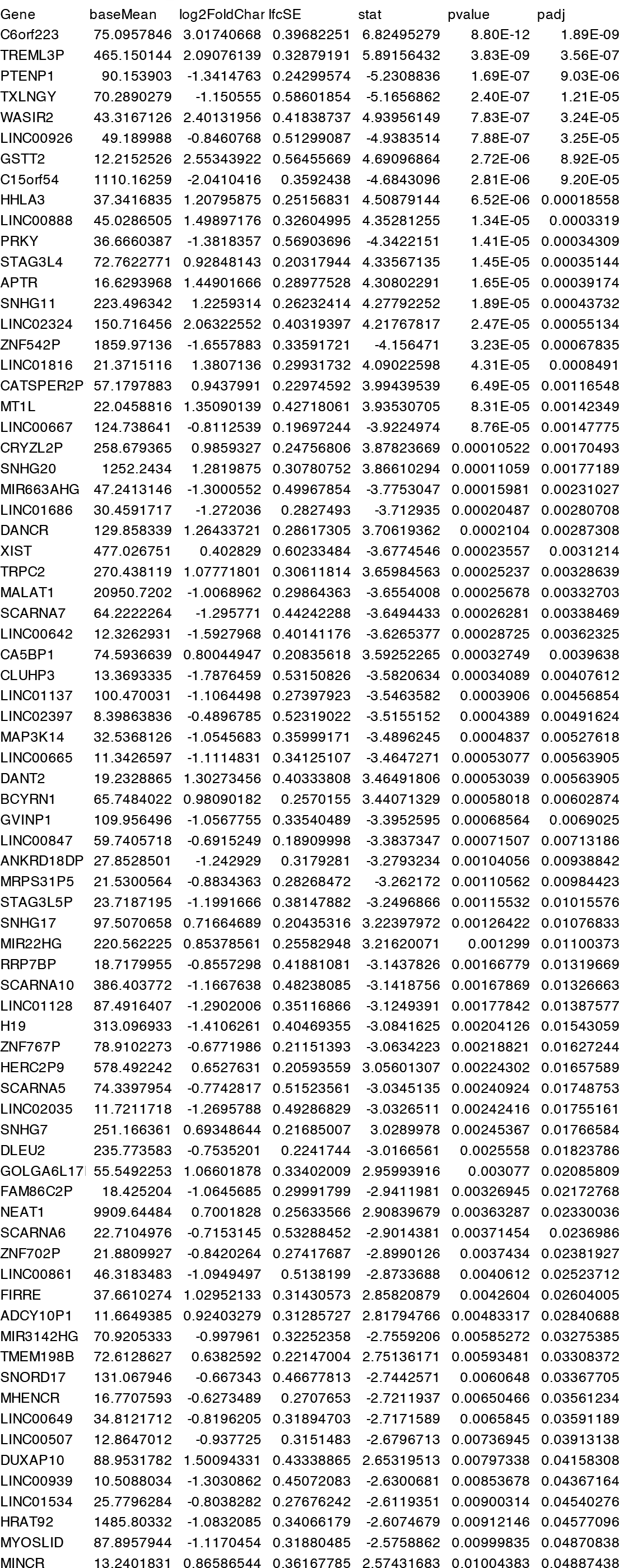
Non-coding RNA ET vs CTRL FDR <0.05.

**TableS2B:**
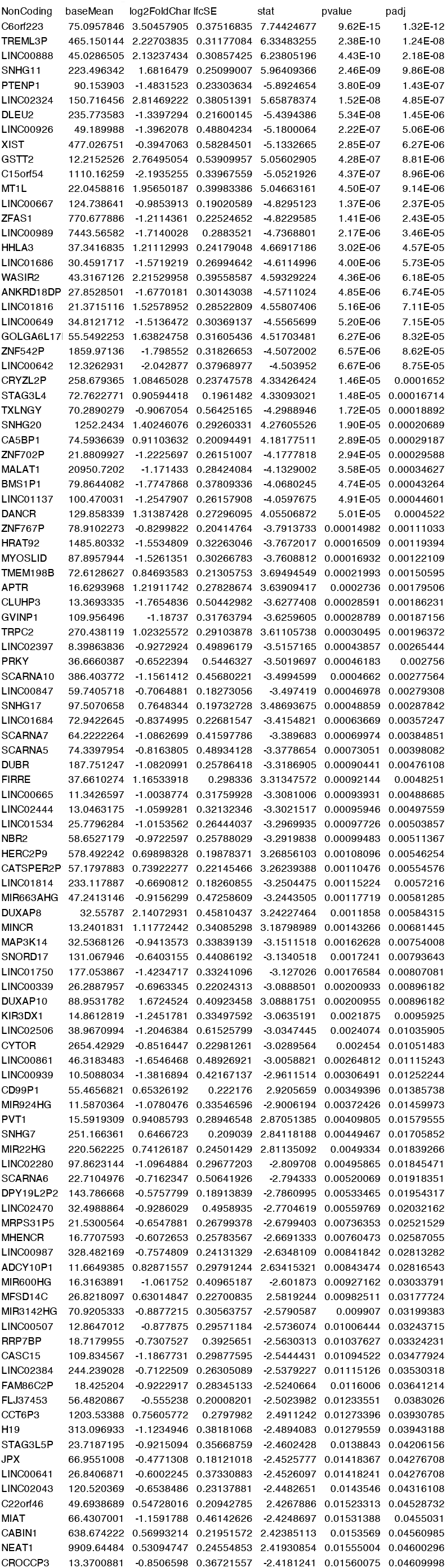
Non-coding RNA PV vs CTRL FDR <0.05.

**TableS2C:**
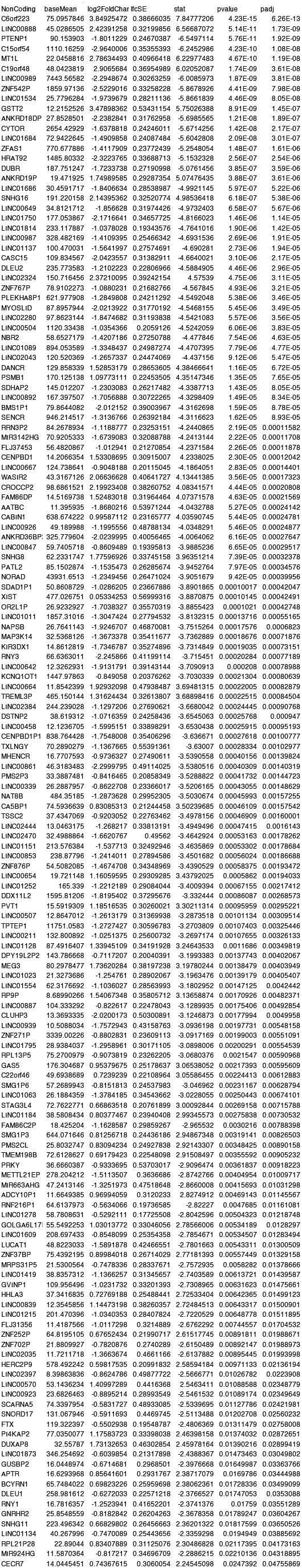
Non-coding RNA MF vs CTRL FDR <0.05.

**Table S3A:** Genes Upregulated in MF and Downregulated in the RUX-treated cohort.

| **Gene** | **Name** |
| --- | --- |
| APLP1 | amyloid beta precursor like protein 1 |
| IFI6 | interferon alpha inducible protein 6 |
| GCH1 | GTP cyclohydrolase 1 |
| CAMK2A | calcium/calmodulin dependent protein kinase II alpha |
| TNIP2 | TNFAIP3 interacting protein 2 |
| IFIT1 | interferon induced protein with tetratricopeptide repeats 1 |
| NDRG3 | NDRG family member 3 |
| IFIT2 | interferon induced protein with tetratricopeptide repeats 2 |
| PLEKHM1 | pleckstrin homology and RUN domain containing M1 |
| JOSD1 | Josephin domain containing 1 |
| MYLIP | myosin regulatory light chain interacting protein |
| FAM89B | family with sequence similarity 89 member B |
| SCIN | scinderin |
| NPL | N-acetylneuraminate pyruvate lyase |
| ODF3B | outer dense fiber of sperm tails 3B |
| EIF1AY | eukaryotic translation initiation factor 1A Y-linked |
| IFI27L2 | interferon alpha inducible protein 27 like 2 |
| ISCU | iron-sulfur cluster assembly enzyme |

**Table S3B:** Genes Downreglated in MF and Upregulated n the RUX-treated cohort.

| **Gene** | **Name** |
| --- | --- |
| LINC00926 | long intergenic non-protein coding RNA 926 |
| SEMA3C | semaphorin 3C |
| ADGRG7 | adhesion G protein-coupled receptor G7 |
| CXCR5 | C-X-C motif chemokine receptor 5 |
| LOC101927151 | uncharacterized LOC101927151 |
| SMC5 | structural maintenance of chromosomes 5 |
| PER3 | period circadian regulator 3 |
| SNORD17 | small nucleolar RNA, C/D box 17 |
| LMO7 | LIM domain 7 |

**Table S4A:**
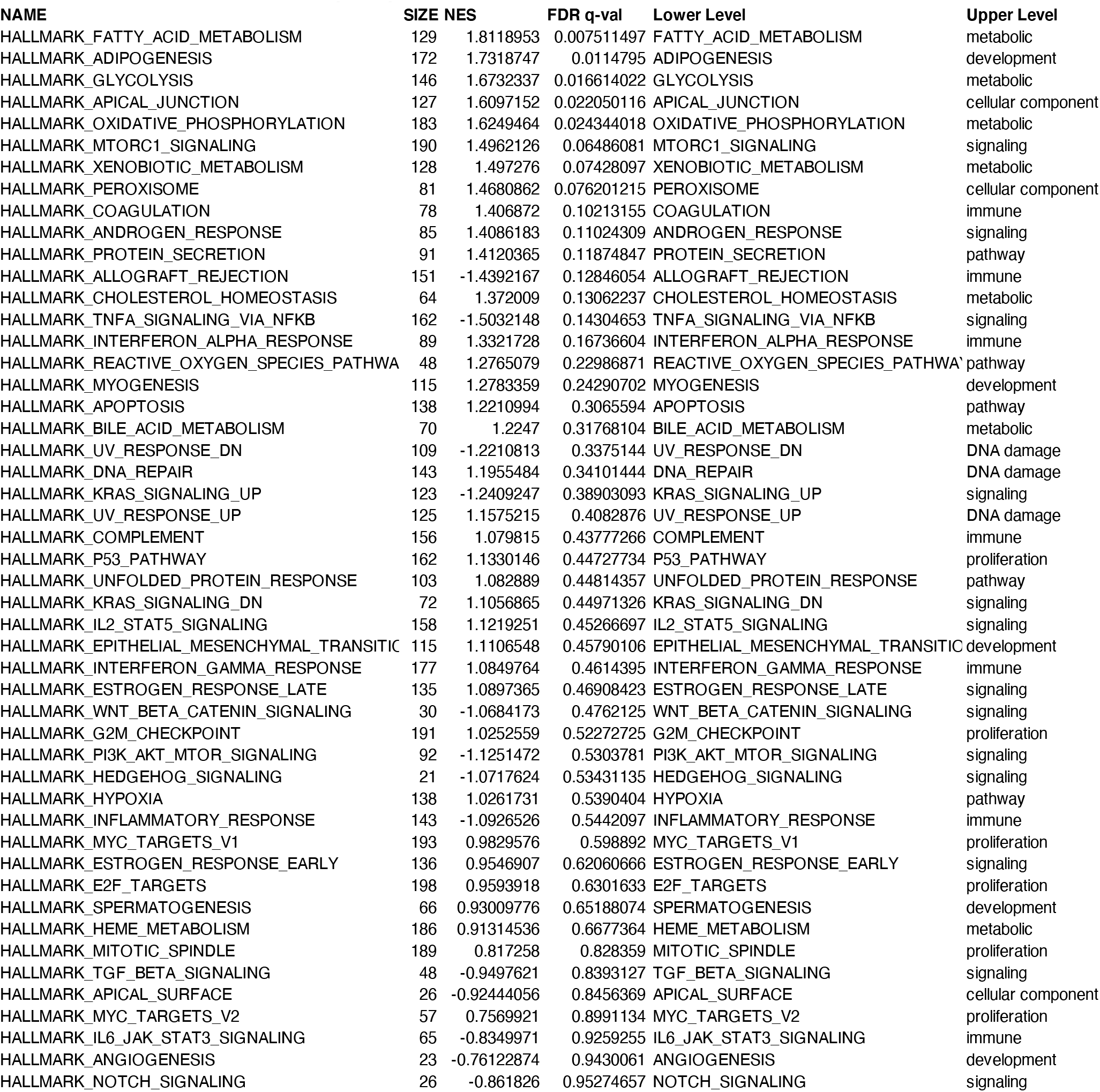
Gene Set Enrichment Pathway Analysis ET vs CTRL.

**Table S4B:** Gene Set Enrichment Pathway Analysis PV vs CTRL.

| NAME | SIZE | NES | FDR q-val | Lower Level | Upper Level |
| --- | --- | --- | --- | --- | --- |
| HALLMARK_ADIPOGENESIS | 172 | 1.7491294 | 0.012118747 | ADIPOGENESIS | development |
| HALLMARK_GLYCOLYSIS | 146 | 1.6530863 | 0.01930149 | GLYCOLYSIS | metabolic |
| HALLMARK_APICAL_JUNCTION | 127 | 1.660242 | 0.025757005 | APICAL_JUNCTION | cellular component |
| HALLMARK_FATTY_ACID_METABOLISM | 129 | 1.6055912 | 0.026302228 | FATTY_ACID_METABOLISM | metabolic |
| HALLMARK_COAGULATION | 78 | 1.5808175 | 0.02970178 | COAGULATION | immune |
| HALLMARK_OXIDATIVE_PHOSPHORYLATION | 183 | 1.5442894 | 0.039326318 | OXIDATIVE_PHOSPHORYLATION | metabolic |
| HALLMARK_CHOLESTEROL_HOMEOSTASIS | 64 | 1.5314032 | 0.04042827 | CHOLESTEROL_HOMEOSTASIS | metabolic |
| HALLMARK_XENOBIOTIC_METABOLISM | 128 | 1.5096257 | 0.042610623 | XENOBIOTIC_METABOLISM | metabolic |
| HALLMARK_INTERFERON_ALPHA_RESPONSE | 89 | 1.518906 | 0.04321455 | INTERFERON_ALPHA_RESPONSE | immune |
| HALLMARK_HEME_METABOLISM | 186 | 1.4567754 | 0.0673379 | HEME_METABOLISM | metabolic |
| HALLMARK_PEROXISOME | 81 | 1.4418689 | 0.07086366 | PEROXISOME | cellular component |
| HALLMARK_MTORC1_SIGNALING | 190 | 1.3795087 | 0.12471676 | MTORC1_SIGNALING | signaling |
| HALLMARK_MYOGENESIS | 115 | 1.3494041 | 0.14121492 | MYOGENESIS | development |
| HALLMARK_EPITHELIAL_MESENCHYMAL_TRANSITION | 115 | 1.356965 | 0.1420532 | EPITHELIAL_MESENCHYMAL_TRANSITION | development |
| HALLMARK_BILE_ACID_METABOLISM | 70 | 1.3197786 | 0.17186424 | BILE_ACID_METABOLISM | metabolic |
| HALLMARK_P53_PATHWAY | 162 | 1.2998974 | 0.17200567 | P53_PATHWAY | proliferation |
| HALLMARK_INTERFERON_GAMMA_RESPONSE | 177 | 1.3008407 | 0.18022385 | INTERFERON_GAMMA_RESPONSE | immune |
| HALLMARK_PROTEIN_SECRETION | 91 | 1.3067324 | 0.18193667 | PROTEIN_SECRETION | pathway |
| HALLMARK_KRAS_SIGNALING_DN | 72 | 1.2802348 | 0.18998314 | KRAS_SIGNALING_DN | signaling |
| HALLMARK_DNA_REPAIR | 143 | 1.2603142 | 0.20102613 | DNA_REPAIR | DNA damage |
| HALLMARK_IL2_STAT5_SIGNALING | 158 | 1.2645485 | 0.20397907 | IL2_STAT5_SIGNALING | signaling |
| HALLMARK_ALLOGRAFT_REJECTION | 151 | -1.3648527 | 0.2617964 | ALLOGRAFT_REJECTION | immune |
| HALLMARK_UV_RESPONSE_UP | 125 | 1.1793895 | 0.34141245 | UV_RESPONSE_UP | DNA damage |
| HALLMARK_COMPLEMENT | 156 | 1.171932 | 0.34395382 | COMPLEMENT | immune |
| HALLMARK_ESTROGEN_RESPONSE_LATE | 135 | 1.1602746 | 0.35328636 | ESTROGEN_RESPONSE_LATE | signaling |
| HALLMARK_HEDGEHOG_SIGNALING | 21 | -1.2357339 | 0.35349664 | HEDGEHOG_SIGNALING | signaling |
| HALLMARK_TNFA_SIGNALING_VIA_NFKB | 162 | -1.1725798 | 0.3754302 | TNFA_SIGNALING_VIA_NFKB | signaling |
| HALLMARK_ESTROGEN_RESPONSE_EARLY | 136 | 1.1259818 | 0.4149885 | ESTROGEN_RESPONSE_EARLY | signaling |
| HALLMARK_REACTIVE_OXYGEN_SPECIES_PATHWAY | 48 | 1.0937103 | 0.475638 | REACTIVE_OXYGEN_SPECIES_PATHWAY | pathway |
| HALLMARK_APOPTOSIS | 138 | 1.0725611 | 0.5131383 | APOPTOSIS | pathway |
| HALLMARK_ANDROGEN_RESPONSE | 85 | 1.061999 | 0.51994646 | ANDROGEN_RESPONSE | signaling |
| HALLMARK_INFLAMMATORY_RESPONSE | 143 | 1.0041803 | 0.65263337 | INFLAMMATORY_RESPONSE | immune |
| HALLMARK_IL6_JAK_STAT3_SIGNALING | 65 | 0.97972196 | 0.6693822 | IL6_JAK_STAT3_SIGNALING | immune |
| HALLMARK_G2M_CHECKPOINT | 191 | 0.98495644 | 0.67889416 | G2M_CHECKPOINT | proliferation |
| HALLMARK_UNFOLDED_PROTEIN_RESPONSE | 103 | 0.9614258 | 0.6946754 | UNFOLDED_PROTEIN_RESPONSE | pathway |
| HALLMARK_HYPOXIA | 138 | 0.94022757 | 0.7245265 | HYPOXIA | pathway |
| HALLMARK_PI3K_AKT_MTOR_SIGNALING | 92 | 0.9294137 | 0.7290277 | PI3K_AKT_MTOR_SIGNALING | signaling |
| HALLMARK_SPERMATOGENESIS | 66 | 0.9083337 | 0.7548022 | SPERMATOGENESIS | development |
| HALLMARK_MITOTIC_SPINDLE | 189 | 0.8941381 | 0.765224 | MITOTIC_SPINDLE | proliferation |
| HALLMARK_UV_RESPONSE_DN | 109 | -1.0125421 | 0.8153306 | UV_RESPONSE_DN | DNA damage |
| HALLMARK_WNT_BETA_CATENIN_SIGNALING | 30 | 0.8177549 | 0.90225345 | WNT_BETA_CATENIN_SIGNALING | signaling |
| HALLMARK_ANGIOGENESIS | 23 | 0.7963376 | 0.91691846 | ANGIOGENESIS | development |
| HALLMARK_KRAS_SIGNALING_UP | 123 | -0.950727 | 0.9232399 | KRAS_SIGNALING_UP | signaling |
| HALLMARK_E2F_TARGETS | 198 | 0.77223194 | 0.9336657 | E2F_TARGETS | proliferation |
| HALLMARK_NOTCH_SIGNALING | 26 | -0.7739246 | 0.9437037 | NOTCH_SIGNALING | signaling |
| HALLMARK_MYC_TARGETS_V1 | 193 | 0.74576426 | 0.94463915 | MYC_TARGETS_V1 | proliferation |
| HALLMARK_MYC_TARGETS_V2 | 57 | 0.6715539 | 0.97912604 | MYC_TARGETS_V2 | proliferation |
| HALLMARK_APICAL_SURFACE | 26 | -0.8297998 | 0.9997721 | APICAL_SURFACE | cellular component |
| HALLMARK_TGF_BETA_SIGNALING | 48 | -0.8463163 | 1 | TGF_BETA_SIGNALING | signaling |

**Table S4C:**
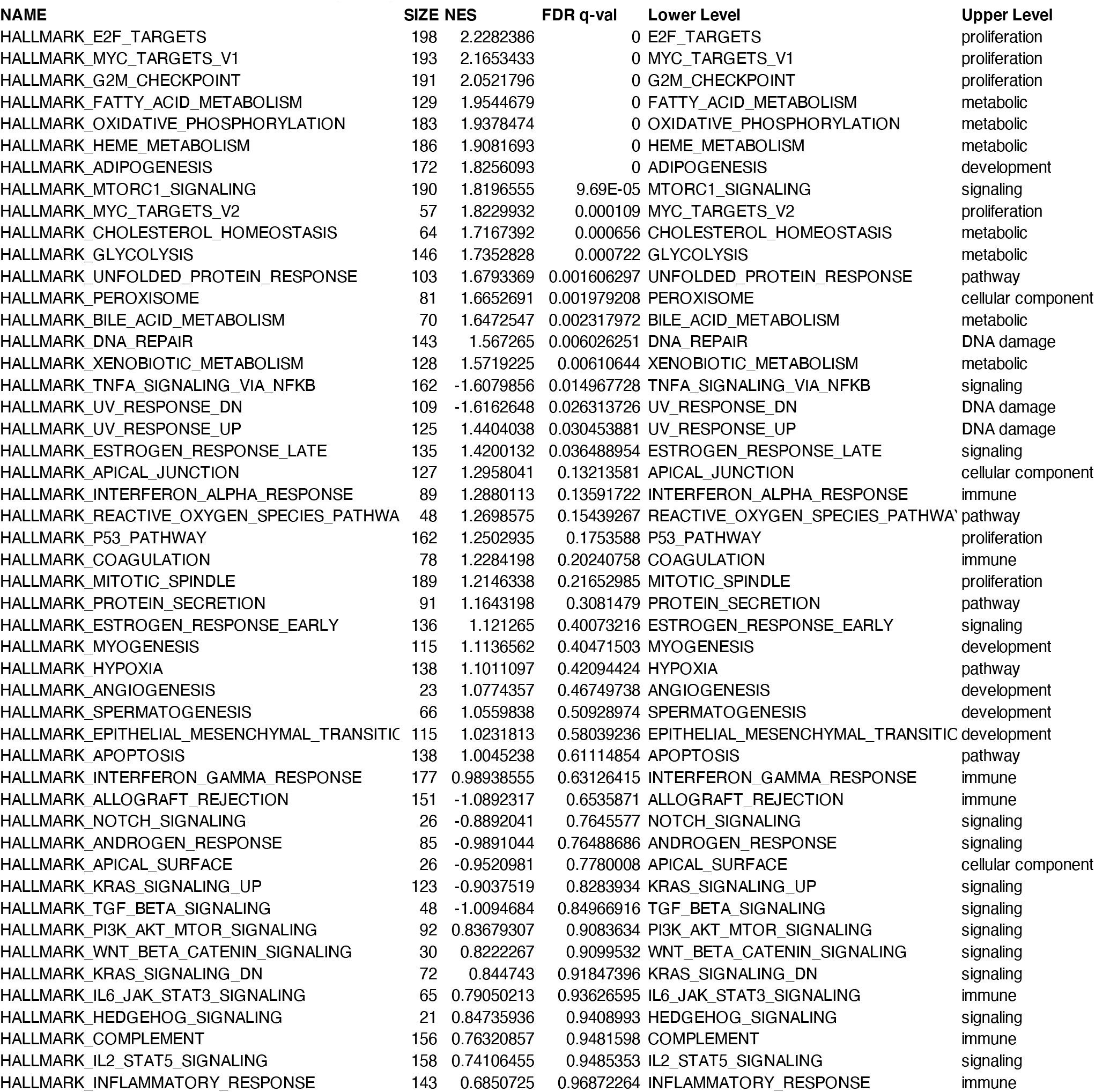
Gene Set Enrichment Pathway Analysis MF vs CTRL.

| NAME | SIZE | NES | FDR q-val | Lower Level | Upper Level |
| --- | --- | --- | --- | --- | --- |
| HALLMARK_E2F_TARGETS | 198 | 2.2282386 | 0 | E2F_TARGETS | proliferation |
| HALLMARK_MYC_TARGETS_V1 | 193 | 2.1653433 | 0 | MYC_TARGETS_V1 | proliferation |
| HALLMARK_G2M_CHECKPOINT | 191 | 2.0521796 | 0 | G2M_CHECKPOINT | proliferation |
| HALLMARK_FATTY_ACID_METABOLISM | 129 | 1.9544679 | 0 | FATTY_ACID_METABOLISM | metabolic |
| HALLMARK_OXIDATIVE_PHOSPHORYLATION | 183 | 1.9378474 | 0 | OXIDATIVE_PHOSPHORYLATION | metabolic |
| HALLMARK_HEME_METABOLISM | 186 | 1.9081693 | 0 | HEME_METABOLISM | metabolic |
| HALLMARK_ADIPOGENESIS | 172 | 1.8256093 | 0 | ADIPOGENESIS | development |
| HALLMARK_MTORC1_SIGNALING | 190 | 1.8196555 | 9.69E-05 | MTORC1_SIGNALING | signaling |
| HALLMARK_MYC_TARGETS_V2 | 57 | 1.8229932 | 0.000109 | MYC_TARGETS_V2 | proliferation |
| HALLMARK_CHOLESTEROL_HOMEOSTASIS | 64 | 1.7167392 | 0.000656 | CHOLESTEROL_HOMEOSTASIS | metabolic |
| HALLMARK_GLYCOLYSIS | 146 | 1.7352828 | 0.000722 | GLYCOLYSIS | metabolic |
| HALLMARK_UNFOLDED_PROTEIN_RESPONSE | 103 | 1.6793369 | 0.001606297 | UNFOLDED_PROTEIN_RESPONSE | pathway |
| HALLMARK_PEROXISOME | 81 | 1.6652691 | 0.001979208 | PEROXISOME | cellular component |
| HALLMARK_BILE_ACID_METABOLISM | 70 | 1.6472547 | 0.002317972 | BILE_ACID_METABOLISM | metabolic |
| HALLMARK_DNA_REPAIR | 143 | 1.567265 | 0.006026251 | DNA_REPAIR | DNA damage |
| HALLMARK_XENOBIOTIC_METABOLISM | 128 | 1.5719225 | 0.00610644 | XENOBIOTIC_METABOLISM | metabolic |
| HALLMARK_TNFA_SIGNALING_VIA_NFKB | 162 | -1.6079856 | 0.014967728 | TNFA_SIGNALING_VIA_NFKB | signaling |
| HALLMARK_UV_RESPONSE_DN | 109 | -1.6162648 | 0.026313726 | UV_RESPONSE_DN | DNA damage |
| HALLMARK_UV_RESPONSE_UP | 125 | 1.4404038 | 0.030453881 | UV_RESPONSE_UP | DNA damage |
| HALLMARK_ESTROGEN_RESPONSE_LATE | 135 | 1.4200132 | 0.036488954 | ESTROGEN_RESPONSE_LATE | signaling |
| HALLMARK_APICAL_JUNCTION | 127 | 1.2958041 | 0.13213581 | APICAL_JUNCTION | cellular component |
| HALLMARK_INTERFERON_ALPHA_RESPONSE | 89 | 1.2880113 | 0.13591722 | INTERFERON_ALPHA_RESPONSE | immune |
| HALLMARK_REACTIVE_OXYGEN_SPECIES_PATHWA | 48 | 1.2698575 | 0.15439267 | REACTIVE_OXYGEN_SPECIES_PATHWA | pathway |
| HALLMARK_P53_PATHWAY | 162 | 1.2502935 | 0.1753588 | P53_PATHWAY | proliferation |
| HALLMARK_COAGULATION | 78 | 1.2284198 | 0.20240758 | COAGULATION | immune |
| HALLMARK_MITOTIC_SPINDLE | 189 | 1.2146338 | 0.21652985 | MITOTIC_SPINDLE | proliferation |
| HALLMARK_PROTEIN_SECRETION | 91 | 1.1643198 | 0.3081479 | PROTEIN_SECRETION | pathway |
| HALLMARK_ESTROGEN_RESPONSE_EARLY | 136 | 1.121265 | 0.40073216 | ESTROGEN_RESPONSE_EARLY | signaling |
| HALLMARK_MYOGENESIS | 115 | 1.1136562 | 0.40471503 | MYOGENESIS | development |
| HALLMARK_HYPOXIA | 138 | 1.1011097 | 0.42094424 | HYPOXIA | pathway |
| HALLMARK_ANGIOGENESIS | 23 | 1.0774357 | 0.46749738 | ANGIOGENESIS | development |
| HALLMARK_SPERMATOGENESIS | 66 | 1.0559838 | 0.50928974 | SPERMATOGENESIS | development |
| HALLMARK_EPITHELIAL_MESENCHYMAL_TRANSITIC | 115 | 1.0231813 | 0.58039236 | EPITHELIAL_MESENCHYMAL_TRANSITIC | development |
| HALLMARK_APOPTOSIS | 138 | 1.0045238 | 0.61114854 | APOPTOSIS | pathway |
| HALLMARK_INTERFERON_GAMMA_RESPONSE | 177 | 0.98938555 | 0.63126415 | INTERFERON_GAMMA_RESPONSE | immune |
| HALLMARK_ALLOGRAFT_REJECTION | 151 | -1.0892317 | 0.6535871 | ALLOGRAFT_REJECTION | immune |
| HALLMARK_NOTCH_SIGNALING | 26 | -0.8892041 | 0.7645577 | NOTCH_SIGNALING | signaling |
| HALLMARK_ANDROGEN_RESPONSE | 85 | -0.9891044 | 0.76488686 | ANDROGEN_RESPONSE | signaling |
| HALLMARK_APICAL_SURFACE | 26 | -0.9520981 | 0.7780008 | APICAL_SURFACE | cellular component |
| HALLMARK_KRAS_SIGNALING_UP | 123 | -0.9037519 | 0.8283934 | KRAS_SIGNALING_UP | signaling |
| HALLMARK_TGF_BETA_SIGNALING | 48 | -1.0094684 | 0.84966916 | TGF_BETA_SIGNALING | signaling |
| HALLMARK_PI3K_AKT_MTOR_SIGNALING | 92 | 0.83679307 | 0.9083634 | PI3K_AKT_MTOR_SIGNALING | signaling |
| HALLMARK_WNT_BETA_CATENIN_SIGNALING | 30 | 0.8222267 | 0.9099532 | WNT_BETA_CATENIN_SIGNALING | signaling |
| HALLMARK_KRAS_SIGNALING_DN | 72 | 0.844743 | 0.91847396 | KRAS_SIGNALING_DN | signaling |
| HALLMARK_IL6_JAK_STAT3_SIGNALING | 65 | 0.79050213 | 0.93626595 | IL6_JAK_STAT3_SIGNALING | immune |
| HALLMARK_HEDGEHOG_SIGNALING | 21 | 0.84735936 | 0.9408993 | HEDGEHOG_SIGNALING | signaling |
| HALLMARK_COMPLEMENT | 156 | 0.76320857 | 0.9481598 | COMPLEMENT | immune |
| HALLMARK_IL2_STAT5_SIGNALING | 158 | 0.74106455 | 0.9485353 | IL2_STAT5_SIGNALING | signaling |
| HALLMARK_INFLAMMATORY_RESPONSE | 143 | 0.6850725 | 0.96872264 | INFLAMMATORY_RESPONSE | immune |

**Table S5A:**
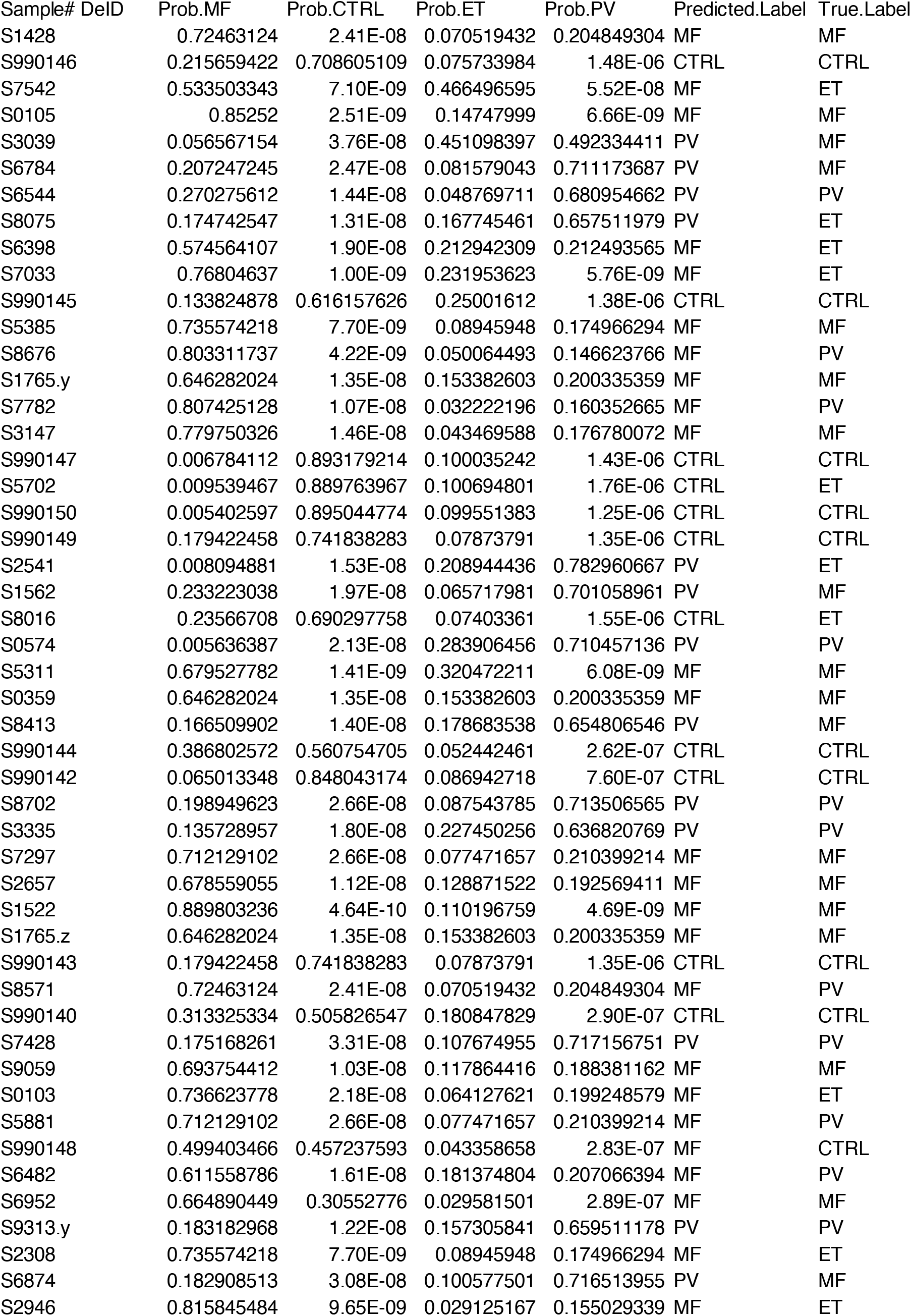
Predicted Probabilities of Lasso Base Model (Age, Gender, Mutation Status)

**Table S5B:**
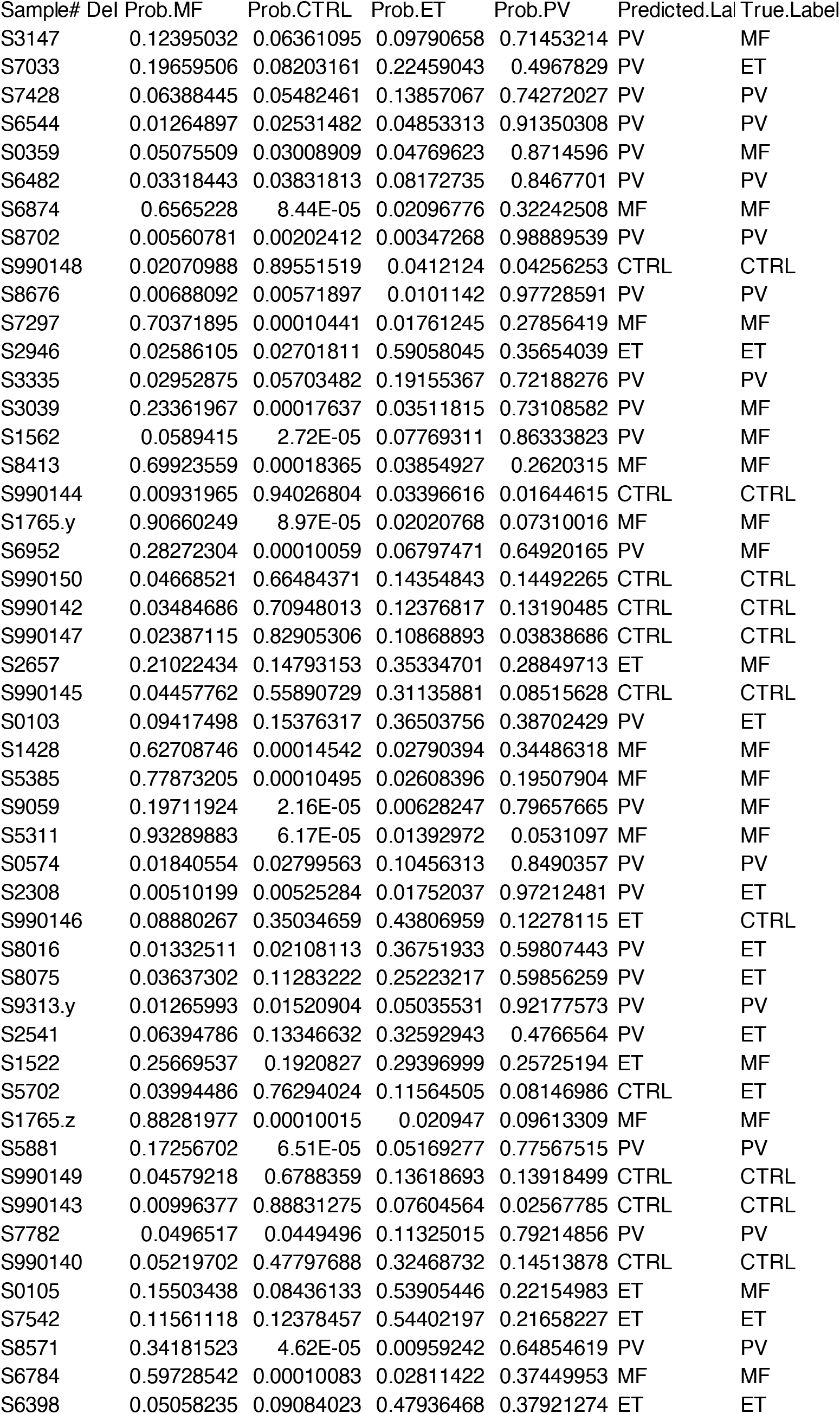
Predicted Probabilities of Lasso Model with the Entire Transcriptome.

**Table S5C:**
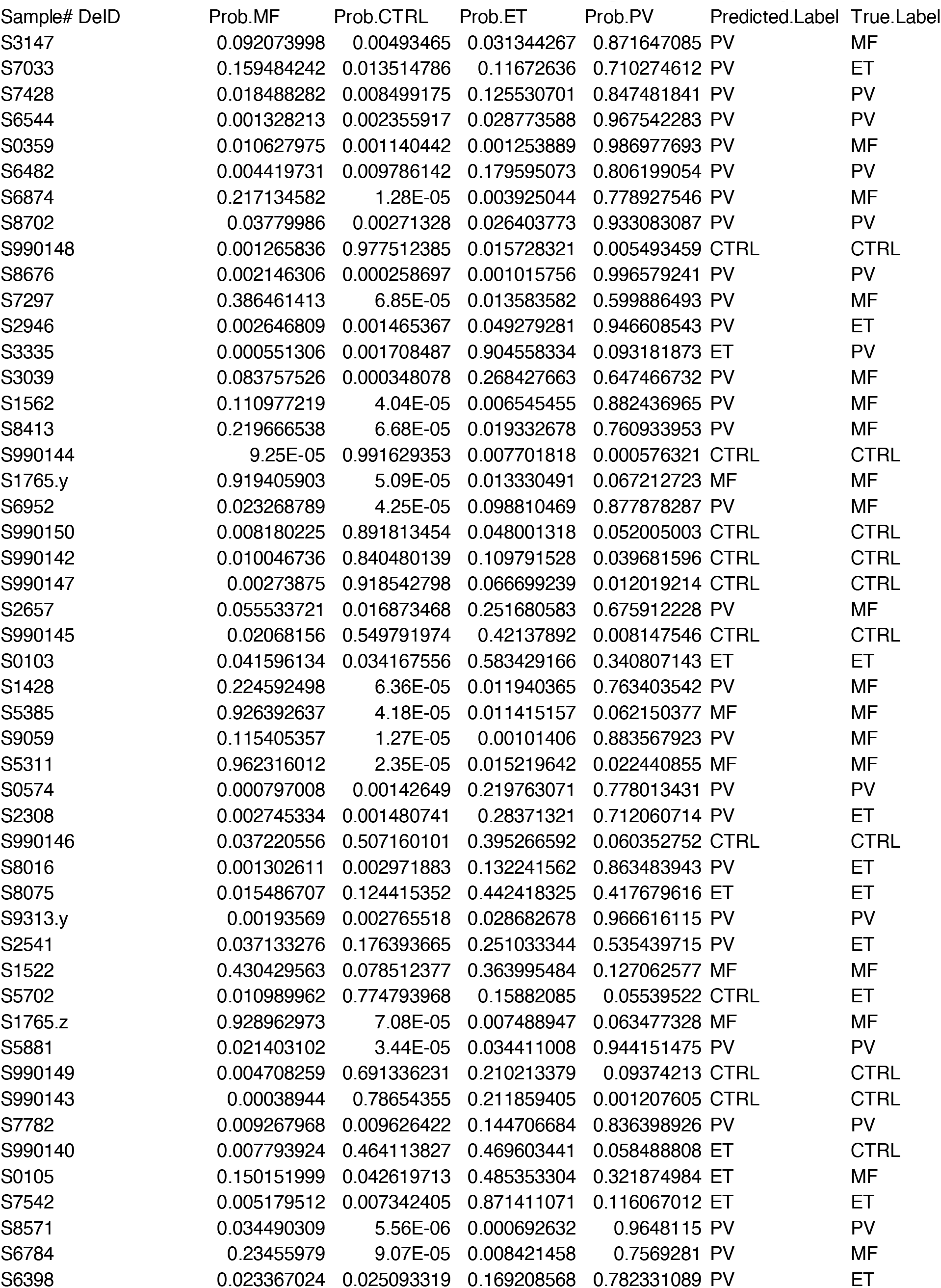
Predicted Probabilities of Lasso Model with the Progressive Transcriptome.

